# Ephemeral resources promote adaptive sociality in wild guppies

**DOI:** 10.1101/2022.05.20.492799

**Authors:** Lysanne Snijders, Stefan Krause, Alan Novaes Tump, Michael Breuker, Indar W. Ramnarine, Ralf H.J.M. Kurvers, Jens Krause

## Abstract

Sociality can facilitate the discovery of novel resources, yet it can also entail costs, including increased competition, aggression, sexual harassment and disease transmission. Selection should therefore favour social flexibility as a response to changes in resource conditions, allowing individuals to adjust their social associations optimally. Quantifying these social adjustments and how they allow individuals to benefit in dynamically changing resource environments can provide insight into the adaptive capacity of wild animal populations. We experimentally studied social flexibility in wild guppies (*Poecilia reticulata*) by examining their fission-fusion dynamics. Using mixed-sex sets of individually marked fish, we manipulated recent foraging experience by presenting them with unpredictable, ephemeral food patches or control patches in the wild. Guppies exposed to ephemeral food patches significantly increased their sociality, doubling the proportion of time spent social compared to controls. This shift was driven by an increase in the probability of starting social contact and a decrease in the probability of ending social contact. Stronger increases in sociality were associated with individuals more often locating ephemeral food patches, mediated by foraging motivation. Importantly, increased sociality was not merely the result of local aggregation, as exposure to ephemeral food patches increased, rather than reduced, spatial activity. And stronger spatial activity did not result in more time spent social. In a second experiment, fish were briefly presented with food patches they could see and smell but not actually consume. Again, fish increased their level of sociality compared to control, indicating that the observed social response was not merely driven by satiation-related changes in activity budgets. Together, our results highlight the adaptive and dynamic nature of fission-fusion social systems and support the view that wild animals adaptively shape their social environment in response to fluctuations in resource variability.

## 1. Introduction

The selective advantages of living in groups have shaped the evolution of sociality across numerous taxa (Garcia and De Monte, 2013; Rubenstein and Abbot, 2017). Indeed, social attributes positively predict fitness measures in a variety of species (Formica et al., 2012; Ryder et al., 2009; Silk et al., 2003). The fitness advantages of sociality can emerge through various pathways (Krause and Ruxton, 2002), such as facilitating predation avoidance, mating opportunities, or foraging success (Karplus et al., 2006; Oh and Badyaev, 2010; Shier and Owings, 2007; Snijders et al., 2021). An increase in foraging success can be especially impactful because it can positively affect both survival and reproduction (Abrahams, 1993; Blanckenhorn, 1991; Clark and Mangel, 1986; Clay et al., 2018; Ellis et al., 2017; Morse and Stephens, 1996).

Sociality facilitates resource discovery in many animals (Galef and Giraldeau, 2001; Oestreich et al., 2026). Specifically, social relationships can help locate novel resources (Firth et al., 2016; Hasenjager et al., 2020a, 2020b; Heinen et al., 2022), with individuals that spend more time socially or are more centrally positioned in the social network typically discovering resources more quickly and frequently (Aplin et al., 2012; Carter et al., 2016; Snijders et al., 2019, 2018) or experiencing less food intake variance (Jolles et al., 2017). A variety of mechanisms can underlie this positive social effect on novel resource localisation, including social facilitation, i.e., the mere presence or behaviour of others increases the probability of a known behaviour, such as exploration or foraging (Dindo et al., 2009) and local enhancement, i.e., attraction to a location because others are present (Avarguès-Weber and Chittka, 2014; Webster and Laland, 2012). Increased sociality can, however, also be costly, e.g., by increasing the risk of competition, (sexual) harassment, and disease exposure (Darden et al., 2009; Godfrey et al., 2009; Leu et al., 2016). Animals are therefore expected to flexibly adapt their social state to prevailing resource conditions (Fisher et al., 2021; Komdeur and Ma, 2021).

When the prevailing resource conditions make social associations less costly or more beneficial, e.g., by reducing competition or facilitating resource localisation, sociality should increase. In support, various field studies have found that social behaviour often varies or changes with natural resource variability (Brown, 1988; Chapman et al., 1995; Dorning and Harris, 2019; Egert-Berg et al., 2018; Foster et al., 2012). Furthermore, experimental evidence suggests that individuals may strategically adjust their social behaviour in response to changes in foraging opportunities (Grubb, 1987; Kulahci et al., 2018). For example, an experimental field study showed that without easy access to supplemental food, wild blue tits (*Cyanistes caeruleus*) and great tits (*Parus major*) increasingly foraged in multispecies flocks (Grubb, 1987), presumably to locate alternative feeding opportunities more effectively. Such changes in sociality in response to changes in resource variability likely result in foraging benefits, but these benefits often remain unquantified. Quantifying the speed and magnitude of social adjustment and testing if such adjustments are indeed beneficial in fitness-relevant contexts is crucial to assess if animals can mitigate resource fluctuations, or other forms of rapid environmental change, through social flexibility (i.e., ’social buffering’), allowing us to better understand the adaptive capacity of social systems.

Here, we investigated whether changes in resource availability induce strategic adjustments in the sociality of wild Trinidadian guppies (*Poecilia reticulata*) and whether stronger social adjustment indeed results in larger foraging benefits. Trinidadian guppies are an ideal study model for this question, since earlier field research showed that guppies exhibit fission-fusion dynamics, switching frequently between being alone and being in close proximity to conspecifics (Wilson et al., 2015, 2014). Individual guppies consistently vary in how much time they spend social (Krause et al., 2017), and this ’social time’ positively correlates with their individual success in locating ephemeral food patches (Snijders et al., 2019, 2018). Moreover, a recent field study manipulated the number of guppies in a given pool, showing that an individual’s food patch discovery and food consumption increased with the number of conspecifics (Snijders et al., 2021). Being more social thus has a clear positive effect on individual foraging success, likely through increased exposure to social information in the form of visual and olfactory cues. Indeed, guppies have been a key model species for the study of social foraging and social learning of foraging information (Clément et al., 2017; Day et al., 2001; Franks and Marshall, 2013; Hasenjager and Dugatkin, 2017; Lachlan et al., 1998; Laland and Williams, 1997; Morrell et al., 2008; Reader et al., 2003; Swaney et al., 2001), yet whether wild guppies strategically alter their social state and so effectively increase foraging efficiency in response to fluctuating resource availability remains to be tested.

Building on these insights, we experimentally studied the effects of ephemeral food patch availability on the social dynamics and foraging success of 18 mixed-sex sets of wild guppies in natural rainforest pools. For readability purposes, we refer to these sets as ’groups’ but emphasise that this terminology is not meant to indicate that these fish primarily move around as cohesive units. Instead, these fish show fission-fusion dynamics during which they frequently alternate between solitary and non-solitary (i.e., in proximity of at least one other group member) states. Using a counter-balanced within-group design, we exposed each group, *in situ*, to two series of food and control presentations, while observing their social dynamics before, in between and after. First, we tested whether groups were repeatable in the overall time their members spent social under control conditions. Second, we tested whether this time spent social changed following experience with ephemeral food patch availability, predicting that fish would become more social to increase access to social information and better locate patchy, ephemeral food resources (Oestreich et al., 2026). To examine alternative explanations of changes in sociality, such as carry-over of local aggregation or increased spatial activity, we performed several supportive data analyses. Third, we tested whether changes in the time groups spent social correlated with higher individual foraging success. Lastly, in a follow-up experiment, we replaced the consumable food patches with food odour patches and tested if time spent social also changed when fish could perceive ephemeral food patches but not consume them. We designed this second experiment to investigate the potential role of satiation-related changes in activity budgets as the driver of observed changes in sociality (Abrahams, 1993; Borgeaud et al., 2021; Gareta García et al., 2021; Kolluru and Grether, 2005).

## 2. Materials and Methods

### Study area and subjects

The study took place in the rainforest region of the Upper Turure River (10°41’8” N, 61°10’22” W) of Trinidad’s Northern Range. This region is considered a low-predation and resource-poor site (Barbosa et al., 2018; Deacon et al., 2018; Grether et al., 2001). We conducted two experiments: Experiment (Exp) 1 (6 – 24 March 2019) studied how fish adjusted their sociality when experiencing a rapid influx of ephemeral food patches, while Exp 2 (8 – 15 April 2023) studied how fish adjusted their sociality when experiencing a similar setup but without the possibility to become satiated by using food odour patches. We used four (Exp 1) or three (Exp 2) natural pools (approximate surface area range of the pools: 2–5 m^2^; depth range: 5–30 cm), situated at the edge of the stream, which were solidified around the borders to allow a constant in- and outflow of water from the main stream while minimising opportunities for fish migration. The primary substrate of the pools comprised a characteristic mix of gravel, leaves and rock (no plants). We did not measure temperature or oxygen levels. Still, by using natural pools we ensured we would study the guppies in an ecologically relevant environment, including corresponding abiotic conditions, variations in water levels and low (but not absent) predation risk levels. Moreover, guppies could maintain their natural baseline foraging behaviour by allowing exploration and exploitation of low-quality food resources such as algae and detritus (Grether et al., 2001; Zandonà et al., 2011). Guppies originally occurring in these pools were caught and moved to other parts of the stream. For our experimental groups, we collected guppies from a nearby stretch of the river. Upon capture, adult fish (Exp 1: N = 144; Exp 2: N = 160; 50/50 sex-ratio) were measured (Exp 1: females: (Mean ± SD) 24.3 ± 2.4 mm, males: 22.1 ± 1.6 mm; Exp 2: females: 24.5 ± 2.5 mm; males = 23.7 ± 1.6 mm) and individually marked using an established technique of Visible Implant Elastomer (VIE) tagging (©Northwest Marine Technology Inc.) (Croft et al., 2003; Jungwirth et al., 2019; Snijders et al., 2018, 2021). Colour marks were green, red, white or pink and placed either at the relative front or back of an individual. An earlier study with wild guppies showed that the use of red elastomers (which may most closely resemble orange, a relevant colour in guppy mate choice) did not affect the time a male (or a female) spent socially or how it distributed its contact moments (Snijders et al., 2019).

The experimental groups consisted of eight fish (of equal sex ratio), and fish within the same group were caught from the same natural site to retain familiarity. We released the groups in their respective experimental pools and left them to settle overnight, with experimental trials starting the following morning. In Exp 1, four groups lost one fish overnight, resulting in 14 groups of eight fish and four groups of seven (N = 18 groups, **Table 1**). In Exp 2, three groups lost one fish overnight, and two groups lost two fish, resulting in 13 groups of eight, three groups of seven, and two groups of six fish (N = 18 groups, **Table 2**). In addition, two groups in Exp 2 were completely replaced after losing three or more fish, likely due to overnight flooding. In Exp 1, the number of groups studied per unique pool ranged between three and six, while in Exp 2, there were six groups studied per unique pool.

**Table 1.** Schematic overview of the design for Experiment 1. Groups of seven or eight individuals, distributed across four pools, received the food patch treatment on day 1 and control on day 2, or vice versa. Within each day, we observed social dynamics using focal observations before (S0), between (S1), and after (S2) the two patch presentation series. The series included presentations of either ephemeral food patches (F1 & F2) or control patches (C1 & C2).

| Day 1 |  |  |  |  | Day 2 |  |  |  |  | Groups | Individuals |
| --- | --- | --- | --- | --- | --- | --- | --- | --- | --- | --- | --- |
| Food patches |  |  |  |  | Control patch |  |  |  |  | N = 9 | N = 71 |
| S0 | F1 | S1 | F2 | S2 | S0 | C1 | S1 | C2 | S2 |  |  |
| Control patches |  |  |  |  | Food patches |  |  |  |  | N = 9 | N = 69 |
| S0 | C1 | S1 | C2 | S2 | S0 | F1 | S1 | F2 | S2 |  |  |

Our research complied with the law and relevant ethical regulations at the time of and in the country of study (Trinidad and Tobago). Specifically, the study was performed in accordance with the ‘Basic Principles Governing the Use of Live Animals and Endangered Species in Research’ at the University of the West Indies as part of the ‘Policy and Procedures on Research Ethics’ of the University of the West Indies Campus Research Ethics Committee, St. Augustine (Ref: CREC-SA.270/03/2023). We released all subjects on the day of their final trials.

### Experiment 1

#### Experimental design

Exp 1 used a counter-balanced within-group design in which every group experienced both a food patch treatment and a procedural control on two consecutive days, with half of the groups receiving the control first and the other half receiving the treatment first (**Table 1**). Each experimental day consisted of three focal observation sessions (S0, S1 and S2; approx. 25-30 min each), separated by two series of 60 patch presentations (F1 and F2 or C1 and C2; 30 min each). Focal observations quantified social dynamics before, between and after the presentations, whereas patch presentations quantified individual foraging success by determining whether or not individuals arrived at an ephemeral food patch. Session S0 served to quantify baseline levels of social dynamics, while sessions S1 and S2 served to quantify social dynamics under either ephemeral food or procedural control conditions. From here onwards we refer to these focal observation sessions of social dynamics as ‘social sessions’. An experimental day thus comprised: social session (S0), patch presentation series (F1 or C1), two-minute break, social session (S1), larger break (30-45 minutes), patch presentation series (F2 or C2), two-minute break, social session (S2) (Supporting information **Fig. S1**).

#### Focal observations of social dynamics

Each experimental day, we performed focal follow-observation sessions (‘social sessions’) of each group member before (S0), between (S1), and after (S2) the food or control presentations (**Table 1**, Supporting information **Fig. S1**). Each individual was followed for three minutes during each session, and we noted every ten seconds whether the focal fish was alone or social. Staying consistent with previous observation protocols, we considered a guppy social when it was within four body lengths of another guppy (Krause et al., 2017; Snijders et al., 2019; Wilson et al., 2014). Ten seconds was considered an appropriate time interval, since earlier research (in a denser population) showed that guppies on average encounter other individuals, or shoals, every fourteen seconds (Croft et al., 2003). The naturally clear and shallow pools allowed us to visually track the fish continuously. The observer sat quietly next to the pool in a location where they could oversee the entire pool with minimum glare or other obstruction. Guppies rarely went into hiding but occasionally went out of sight, for example when foraging under a leaf. If no other guppies were observed entering or exiting the same location, we would assume the focal individual was alone. If, on rare occasions, we lost track of a fish for more than 20 seconds, we would repeat the entire observation. Fish primarily swam just above the bottom of the pool, allowing us to also reliably note their spatial zone in the pool every ten seconds. Spatial zones were identified by determining the nearest of five (a-priori assigned) patch presentation locations. These focal observations served as the primary measure of social and spatial dynamics and were later summarised into social (or spatial) transition probabilities and time spent social (see ‘Statistical analyses’). In total, we performed 108 social observation sessions (six social sessions per group).

During the focal observations, we additionally continuously scored events of sigmoidal display (an S-shaped body posture shown by males to display their body to females), sexual harassment (active chasing and attempts to make physical contact performed by males towards females) and general aggression (active chasing and attempts to make physical contact, excluding sexual harassment) performed or received by the focal fish. To minimise selection bias, the focal observation order was determined haphazardly by a second person (who was not currently observing the fish). Because live observation was required, the focal observations could not be conducted blindly. We acknowledge that this limitation could have introduced an observation bias, but given the magnitude of the effect (see ‘Results’), we believe it is unlikely that this limitation qualitatively influenced our results.

#### Ephemeral patch presentations and foraging observations

Each experimental day, we conducted two series of either food (F1 and F2) or control (C1 and C2) patch presentations (**Table 1**, Supporting information **Fig S1**). One series comprised 60 presentations, simulating an influx of patchy and ephemeral food resources, e.g., fallen fruits or insects in rainforest streams. A series of presentations lasted 30 minutes in total, with a single presentation and a break between two presentations each lasting 15 seconds. Patches were presented at five a-priori assigned locations, roughly evenly distributed throughout the entire pool (with a minimum distance of 50 cm between nearest locations) and marked by naturally occurring white pebbles. The order of presentation location was determined randomly (using a random number generator), with the restriction that within every five trials, each location received a presentation. As a food patch, we used small lead balls (8 mm diameter), which were covered with a mix of gelatine and crushed fish food flakes, including carotenoids (TetraPro©; Spectrum Brands Inc). Food patches were shareable, as multiple individuals could (and would) feed simultaneously. As a control, we used the same balls but without a food cover. During a presentation, while positioned next to the pool, we lowered the ball, which was attached with a monofilament line to a stick, into the water until approximately two cm above the bottom of the pool.

To quantify individual foraging success, we conducted behaviour observations during the patch presentations by scoring the identity and the order of the individuals that arrived within two body lengths of a patch, and, specifically for the ephemeral food treatment, whether they took at least one bite from the patch (referred to as a ‘feeding event’ from here onwards). Feeding events could not be reliably determined during six of the presentations. In addition, during one food presentation series, the food patch was completely depleted before the final presentation and therefore ended 17 presentations early (i.e., 43 presentations instead of 60). In total, we conducted 4,303 successful patch presentations, resulting in 33,481 ephemeral patch arrival (yes/no) observations at the individual level.

### Experiment 2

#### Experimental design

To investigate whether simple energetic changes following food consumption caused most of the observed changes in social dynamics, we performed a complementary second experiment in a subsequent field season. In Exp 2, we replaced the series of food patch presentations with a series of food odour patch presentations. This experiment followed a counter-balanced within-group design, similar to Exp 1. However, in Exp 2, groups received treatment and procedural control presentations on the same day (**Table 2**). This design helped minimise the overnight loss of fish (e.g. due to escape, flooding or predation), which occurred more frequently this field season. An experimental day consisted of two food odour (F1 and F2) and two control (C1 and C2) patch presentation series (7.5 min/series), with each series including nine patch presentations. The odour presentation series were, again, interspersed with focal observation sessions of social dynamics (‘social sessions’; approx. 20-25 min per session). However, to allow each group to be tested within a day, we did not conduct a social session before the first presentation series (S0), deeming the comparison between treatment and procedural control using the remaining social sessions (S1 and S2) sufficient. Specifically, an experimental day consisted of: patch presentations (F1 or C1), a two-minute break, social session (S1), patch presentations (F2 or C2), a two-minute break, social session (S2), a treatment switch break (30-45 minutes), patch presentations (C1 or F1), a two-minute break, social session (S1), patch presentations (C2 or F2), a two-minute break, and a final social session (S2) (Supporting information **Fig. S1**). During Exp 2, before the start of the experimental trials, guppies in all groups were allowed to take a few bites of the food item for a maximum of 1 minute to ensure they would associate the odour food patch with a food resource during the experiments. We left at least 15 minutes between this brief food exposure and the experimental trials.

**Table 2.**
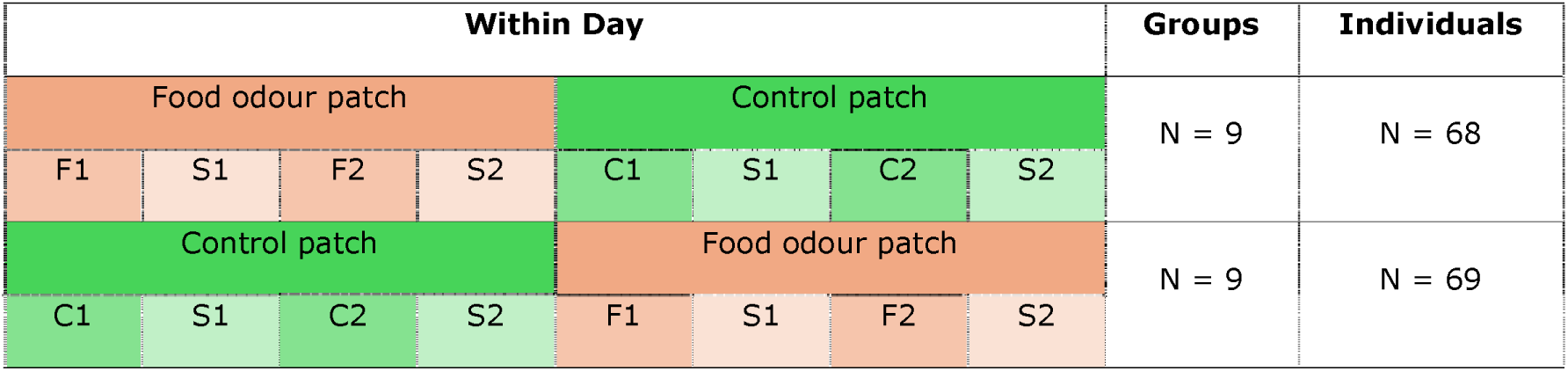
Schematic overview of the design for Experiment 2. Groups of six to eight individuals, distributed across three pools, received food odour patches (F1 & F2) and then control patches (C1 & C2) on the same day, or vice versa. We observed social dynamics during focal observation sessions in between (S1) and after (S2) the two patch presentation series.

#### Focal observations of social dynamics

The focal observations to quantify and compare the social dynamics followed the same observation protocol as in Exp 1, but with no social session before the patch presentations (S1 and S2 only; **Table 2**, Supporting information **Fig. S1**) and each individual being observed for two (instead of three) minutes per session.

#### Ephemeral odour patch presentations and foraging observations

In Exp 2, one session of presentations (food odour or control) consisted of nine presentations, each at a unique location within a pool. Note that we increased the number of within-pool locations and reduced the total number of presentations compared to Exp 1 to minimise potential habituation/learning effects (i.e., the guppies learning that the food odour patch does not provide food access and linking this information to locations). A presentation series lasted a total of 7.5 minutes, with every single presentation taking 30 seconds and each break between two presentations lasting 20 seconds. The nine presentation locations were assigned a-priori and roughly evenly spread throughout each pool (with a minimum distance of 25 cm between the nearest locations). The order of presentation location was determined using a random number generator, with the restriction that each location should receive a presentation. We used the same crushed fish food flakes as in Exp 1 to create the food odour patch and small river pebbles to create the control patch. Both odour patch types were wrapped in polyester plankton mesh (Sefar Petex^®^), creating items of approximately 15 by 15 mm. The mesh allowed the odour to diffuse but prevented the fish from accessing the crushed flakes. Multiple individuals could peck at an odour patch simultaneously. As in Exp 1, we lowered the patch, attached with a monofilament line to a stick, into the water until approximately two cm above the bottom of the pool.

To complement our live observations, we successfully filmed all but four of the presentations using camcorders (SONY HDR-PJ530E) on tripods, allowing us to quantify behaviour at the ephemeral odour patches in more detail using BORIS v 8.20 (Friard and Gamba, 2016) (Supporting information). To evaluate the efficacy of our treatment, we scored the number of pecks made at each odour patch. A detected food odour patch received eight times as many pecks as the control patch (and twelve times when considering patches independent of discovery), verifying that the guppies recognised it as a relevant food source.

### Statistical analyses

#### Calculation of social metrics

We quantified social behaviour from the social sessions by estimating the proportion of time individuals spent in close proximity to a conspecific, hereafter referred to as ‘social time’ (Snijders et al., 2019, 2018; Wilson et al., 2014). Social time was calculated using a Markov chain approach. This approach to measuring social dynamics provides more mechanistic insight into social change than simply measuring the proportion of time spent social, since an increase in social time can result from fish initiating social contacts more frequently, maintaining contacts for longer, or a combination of both.

Specifically, each focal observation was translated into a sequence of behavioural states recorded every 10 s, in which an individual was classified as either alone or social (within four body lengths of a conspecific). From these sequences, we estimated two transition probabilities: Pa→s, the probability of starting social contact (transitioning from alone to social), and Ps→a, the probability of ending social contact (transitioning from social to alone). Social time was then calculated as: Pa→s / (Ps→a + Pa→s). Transition probabilities were estimated from the observed frequencies of behavioural state changes. When one transition type was absent, we added one observation to all transition frequencies to avoid estimating unrealistic transition probabilities of zero. Previous studies showed that both individual- and group-level social traits measured in Trinidadian guppies using this approach were repeatable across water surface area manipulations and between-pool translocations (Krause et al., 2017; Wilson et al., 2015).

Because focal observations lasted only 3 min per individual in Exp 1 and 2 min in Exp 2, our design prioritised repeated measurements before, between and after the patch presentations over longer focal observations of individual fish. Consequently, we focused our analyses on group-level social dynamics (Exp 1 and 2) and, in Exp 1, also on sex-specific group-level social dynamics. For each social session, we pooled behavioural state transitions across all focal individuals within a group (or within each sex) to estimate group-level transition probabilities and social time. Because these group-level estimates were independent, no randomisation procedure was required. Social metrics were calculated in Java (OpenJDK version 21.0.7).

#### General modelling approach

We primarily used (generalised) linear mixed models to address four main research questions. For Exp 1: (*i*) Are group differences in social time repeatable under control conditions? (*ii*) Does ephemeral food patch availability increase group-level social time? (*iii*) Does experimentally induced change in group-level social time predict individual foraging success (i.e., arrival at ephemeral food patches)? And for Exp 2: (*iv*) Do ephemeral food-related increases in social time also occur in the absence of food consumption (i.e., no changes in energy budgets)?

All models were fitted in R (R Core Team, 2025) version 4.5.1 in RStudio version 2025.05.1 (© 2009–2025 Posit Software, PBC), using the *lmerTest* package (optimiser = bobyqa) (Kuznetsova et al., 2017). Model families were Gaussian unless specified otherwise. Residual distribution, overdispersion, presence of outliers and quantile deviations were assessed using the ‘simulateResiduals()’ function of the *DHARMa* package (Hartig, 2021). We tested the significance of fixed effects and interactions by comparing models with and without the term of interest using Type III Wald χ² tests (analysis of deviance). For these comparisons, restricted maximum likelihood (REML) was disabled. To facilitate model fitting and comparison of effect sizes, all continuous predictor variables were centred and standardised (mean = 0, SD = 1). Figures were produced using the *ggplot2* package (Wickham, 2016).

The following subsections describe the individual analyses addressing the four main research questions (*i-iv*). In addition, in the Supporting information we summarise all the variables used throughout the analyses and provide an overview of the statistical models, including their specific biological questions, response variables, predictors, and random effects (Supporting information Tables **S1** and **S2**).

##### (i) Repeatability of social time (Exp 1)

We tested the repeatability of group-level social time by constructing mixed models with group identity as a random effect, using the ‘rpt()’ function (1000 bootstraps) of the *rptR* package (Stoffel et al., 2017). We separately calculated the adjusted repeatability of group-level social time for the control (baseline) and food treatment (Supporting information **Table S2: *i*_a**). Repeatability estimates were adjusted for factors inherent to the study design: social session, treatment sequence, and pool identity, as well as group-level biological variables: group size and group-level foraging motivation. This latter variable was included because a low group-level foraging motivation may reflect a fearful state, leading guppies to be more social temporarily. Excluding these group-level variables did not qualitatively change our conclusions, and we therefore report the adjusted repeatability values.

To examine if ephemeral food patch availability induced variation in group-level social time (e.g., by some groups responding more strongly than others), we tested for equality of variances (Levene’s test) between the three social sessions, for both the food treatment and the control treatment (Supporting information **Table S2: *i*_b**).

##### (ii) Ephemeral food patch effects on average social time (Exp 1)

To test if the time spent social increased following ephemeral food patch availability, we focused on the interaction effect of treatment and social session on group-level social time, using a linear mixed model (N = 108 sessions; Supporting information **Table S2: *ii*_a**). In the case of a significant interaction, we performed posthoc contrasts to determine which treatment and session combinations significantly differed from each other, using the *emmeans* package (method = sequential, simple = each, adjust = mvt) (Lenth, 2021). As control variables, we added the main effects of group size, pool identity and the two-way interactions between treatment and treatment sequence and between treatment and group-level foraging motivation. Group identity, nested within pool identity, was included as a random effect to account for repeated measures within groups. To examine whether ephemeral food patch availability affected group-level social time over a prolonged period, we tested whether treatment had a significant effect on the first social (baseline) session (S0) of the next morning (Supporting information **Table S2: *ii*_b**). Due to the limited sample size for this final analysis (N = 18 for S0 on the second day), we used a linear model that included only treatment.

To test whether effects of ephemeral food patch availability may vary between the sexes, we also ran the above-mentioned models for the sex-specific group-level social times (Supporting information **Table S2: *ii*_c**). Moreover, to obtain a more mechanistic understanding of the social changes, we examined the effect of ephemeral food patches on the (group-level) social state transition probabilities underlying social time (i.e., Ps→a and Pa→s; Supporting information **Table S2: *ii*_d**).

#### Ephemeral food patch effects on sexual behaviour frequency (Exp 1)

To test if ephemeral food patches also affected the frequency of sexual behaviours, we calculated the sum of performed and received sexual display and harassment behaviours for each group per social session. We first tested if group-level social time and sexual behaviour frequency were correlated using a Spearman rank test. We then analysed the effects of the interaction between treatment and social session on group-level sexual behaviour using generalised linear mixed models (family = Poisson). Additionally, we analysed the effect of treatment on group-level sexual behaviour during the following morning. These models followed the same model structure as the models reported above except that they also included an observation-level random effect to effectively deal with overdispersion (Supporting information **Table S2: *ii*_e**). The aggression frequency was low (5% of all focal observations) and, therefore, not further explored.

#### Evaluating local aggregation and the effects on social time (Exp 1)

Social time may increase in response to ephemeral food patch availability due to alternative (i.e. non-strategic) reasons, such as carry-over of local aggregation at the food patches. We therefore conducted two additional analyses, which we report in the Supporting Information.

#### Ephemeral food patch effects on spatial activity (Exp 1)

To further examine the potential role of spatial behaviour in more detail, we evaluated whether spatial activity of fish changed in response to ephemeral food patch availability and whether this may have passively resulted in increased sociality levels. Specifically, we first tested if the interaction between treatment and social session affected the probability of transitioning from one spatial zone to another (Supporting information **Table S2: *ii*_h**). Transition probabilities were estimated from the observed frequencies of individuals changing or remaining in a particular spatial zone between consecutive observation points (i.e., ten seconds later). Similar to the social transition probability analyses, we focused these analyses on the group level, meaning that we estimated each transition probability based on the sum of the number of zone changes across all individuals in a group per social session. Subsequently, we tested whether the probability of transitioning between spatial zones affected social time and the underlying social transition probabilities. For this, we ran three linear mixed models, considering only the food treatment, with either social time, probability to start social contact (Pa→s), or probability to end social contact (Ps→a) as the dependent variable (Supporting information **Table S2: *ii*_i**).

##### (iii) Change in social time as predictor of individual foraging success (Exp 1)

Next, we tested if a change in the time a group spends social in response to ephemeral food patch availability correlated with individual foraging success, predicting that increased group-level sociality would facilitate social information transfer about the presence and location of ephemeral food patches. Specifically, we quantified foraging success as individual arrival at a detected food resource, ‘detected’ meaning that at least one guppy in the pool arrived at the resource. We used this metric as the dependent variable in a generalised linear mixed model (family = binomial; N = 9,663; Supporting information **Table S2: *iii*_a**). The higher number of observations per individual for patch arrival, compared to social states, allowed us to analyse foraging success on the individual level, while still including change in social time as a group-level predictor. This ‘social change’ metric was calculated as the difference between group-level social time after the first food session (S1) and before (S0). Here, we assumed that the first few ephemeral food patch presentations would have already triggered the change in social time we measured in session S1. Social time in session S1 (measured within half an hour of F1) thus serves as a proxy for (unbiased) social time during foraging. The model included the two-way interactions between social change and group-level foraging motivation, social change and sex, and social change and food presentation series. In addition, the model included treatment sequence, size (length in mm, centred on sex), group size, and pool identity as the main effects. Group identity nested in pool identity, individual identity nested in group identity, presentation location nested in pool identity and presentation identity (1,237 levels) were included as random effects.

Lastly, to test if arrival at an ephemeral food patch also predicted food intake and thus was indeed a fitness-relevant measure of foraging success, we tested if the proportion of ephemeral patches visited by an individual positively correlated with the total number of feeding events per individual, using Spearman rank correlations (Supporting information **Table S2: *iii*_b**). We ran these analyses for males and females separately.

##### (iv) Ephemeral food odour patch effects on social time (Exp 2)

To test whether an increase in sociality may be driven merely by activity budget changes following food consumption, we tested whether the time spent social by groups changed following the ephemeral availability of a perceived but non-consumable food patch (food odour patch). We analysed the effect of treatment on group-level social time (average of S1+S2) using a linear mixed model (Supporting information **Table S2: *iv***). As control variables, we included the interaction of treatment with treatment sequence, and the main effects of group size and pool identity. Group identity, nested in pool identity, was again included as a random effect to account for repeated measures within groups. Because we used a non-consumable food patch, we could not quantify foraging motivation accurately, and this variable was therefore not included as a control variable.

## 3. Results

### (i) Repeatability of social time (Exp 1)

#### Control conditions

Wild guppy groups showed repeatable differences in group-level social time under baseline conditions (Control S0-S1-S2: R_adj_ = 0.48, SE = 0.16, P = 0.01) and these group differences were relatively stable across the three social sessions (S0: **0.26**, 95% CI = 0.22-0.31; S1: **0.23**, 95% CI= 0.19-0.27; S2: **0.25**, 95% CI = 0.21-0.29; Levene’s test: W_2,51_ = 0.49, P = 0.61; **Fig. 1A & 1D**).

**Figure 1.**
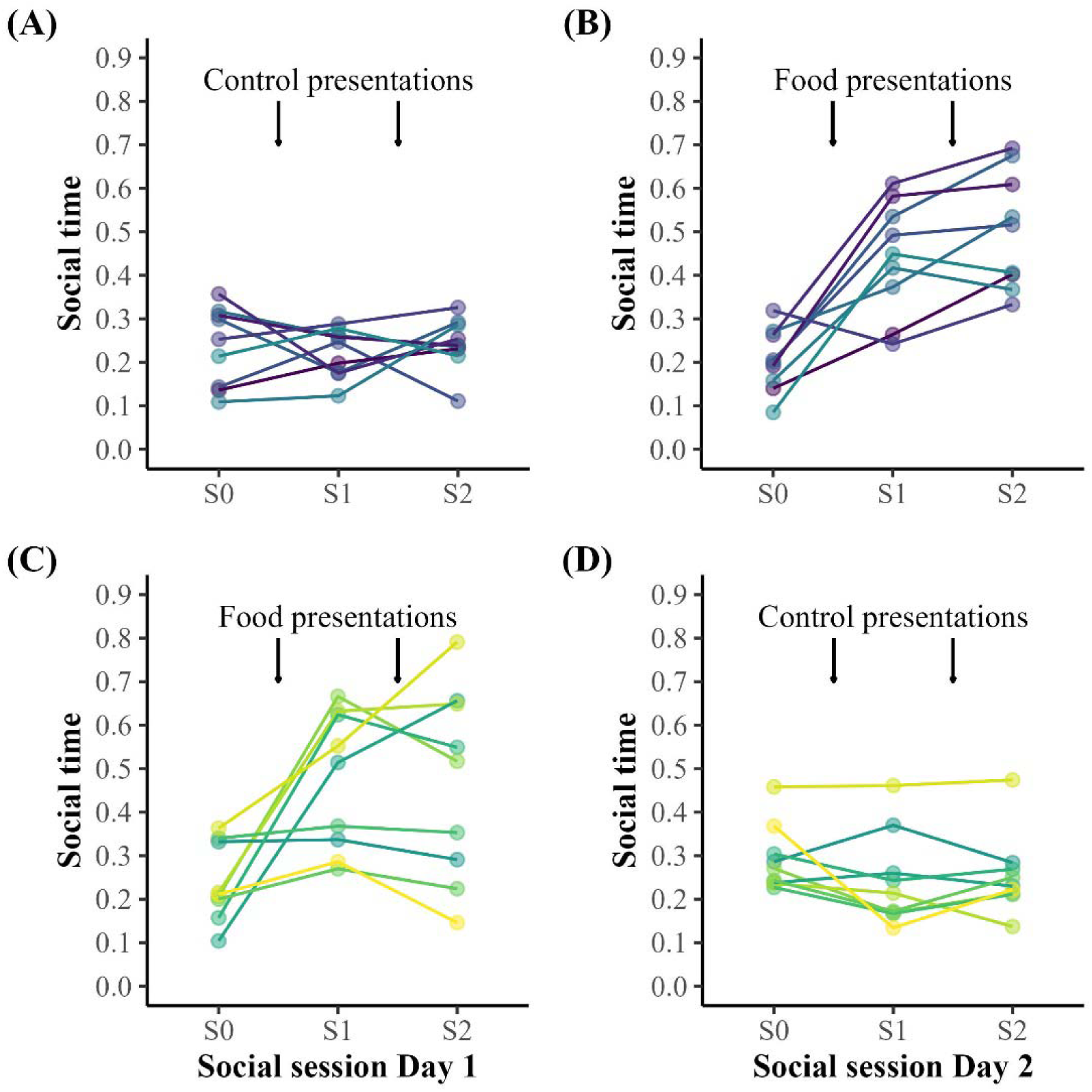
Time spent social by 18 groups of wild guppies before, between, and after two series of repeated food patch or control patch presentations. Arrows indicate the timing of the presentations. A) Groups that received the control patches on the first day, B) received the food patches on the second day. C) Groups that received the food patches on the first day, D) received the control patches on the second day. C & D) The values of the second day’s first social session of social dynamics (S0) visualise the prolonged effects of exposure to ephemeral food patches (during the preceding day)on group-level social time. Colours depict group ID.

#### Ephemeral resource conditions

Repeatability of group-level social time disappeared when comparing social dynamics before and after the initial introduction of ephemeral food patches (Food S0-S1, R_adj_ = 0.00, SE = 0.14, P = 1.00). Moreover, the variance in group-level social time increased (Levene’s test: W_2,51_ = 6.13, P = 0.004, **Fig. 1B & 1C**), suggesting that groups not only socially responded to the ephemeral resources but also that some groups responded more strongly than others. Still, group differences quickly stabilised, indicated by a high level of repeatability when comparing social dynamics after the first and second food patch presentation series (Food S1-S2, R_adj_ = 0.66, SE = 0.18, P = 0.009).

### (ii) Ephemeral food patches positively affect social time (Exp 1)

Recent experience with patchy, ephemeral food resources in the environment significantly increased the average proportion of time spent social (Treatment*Social session: χ2 = 55.65, P < 0.001; Supporting information **Table S3 & S4**, **Fig. 1**), rising twofold (S0: **0.22**, 95% CI = 0.18-0.26; S1: **0.46**, 95% CI= 0.39-0.53; S2: **0.48**, 95% CI = 0.40-0.57). Specifically, more motivated groups changed their social behaviour more strongly than less motivated groups (Treatment*Foraging motivation: χ2 = 9.53, P = 0.002; Supporting information **Table S3**). An increase in social time was still detectable the next morning (Treatment: Estimate (Est) ± SE = 0.09 ± 0.04, N = 18, F = 6.42, P = 0.02, S0 in **Fig. 1D** vs **1B**).

Both females and males were more social following exposure to ephemeral food patches compared to control (Females: Treatment*Social session: χ2 = 27.65, P < 0.001; Supporting information **Table S5**; Males: Treatment*Social session: χ2 = 40.53, P < 0.001; Supporting information **Table S6**). Yet only males still showed an increase in social time during the first social session of the next morning (Females: Treatment: Est ± SE = 0.07 ± 0.06, N = 18, F = 1.54, P = 0.23; Males: Treatment: Est ± SE = 0.11 ± 0.03, N = 18, F = 10.77, P = 0.005).

The time a fish spends social is the result of both the probability of starting social contact and the probability of ending social contact. Analysing these two components of sociality separately, we observed that experience with ephemeral food patches increased the probability of fish starting social contact (Treatment*Social session: χ2 = 28.72, P < 0.001; Supporting information **Table S7**), as well as decreased the probability of fish to end social contact (Treatment*Session: χ2 = 20.59, P < 0.001; Supporting information **Table S8**), suggesting that fish initiated more as well as longer social associations.

#### Ephemeral food patches positively affect sexual behaviours

Sexual behaviour frequency, comprising display and harassment behaviour, increased strongly with group-level social time (r_s_ = 0.54, N = 108, P < 0.001). Mirroring the change in social time, guppies performed twice as many sexual behaviours after experience with ephemeral food patches (Treatment*Social session: χ2 = 22.94, P < 0.001; Supporting information **Table S9 & S10**). Guppies also showed more sexual behaviour in the first social session on the morning after the ephemeral food patch treatment than on the morning after control (Treatment: Est ± SE = 0.87 ± 0.39, N = 18, χ2 = 4.95, P = 0.03).

#### Ephemeral food patch effects on spatial activity

Fish became more likely to transition between spatial zones following experience with patchy, ephemeral resources (Treatment*Social session: χ2 = 7.92, P = 0.02; Supporting information **Table S12 & S13**), indicating increased spatial activity rather than spatial aggregation (S0: **0.17**, 95% CI = 0.14-0.20; S1: **0.23**, 95% CI= 0.19-0.27; S2: **0.22**, 95% CI = 0.19-0.26; Supporting information **Fig. S4**), compared to control. Theoretically, increased spatial activity could, in turn, also lead to a passive increase in social time by increasing the chance of (random) social encounters. However, we found no significant effect of spatial activity on social time (χ2 = 0.10, P = 0.76; Supporting information **Table S14**, **Fig. S5A**). Even though a higher spatial zone transition probability, indeed, predicted a higher probability to start social contact (χ2 = 18.59, P < 0.001; Supporting information **Table S15**, **Fig. S5B**), it also predicted a higher probability to end social contact (χ2 = 4.61, P = 0.03; Supporting information **Table S16**, **Fig. S5C**), leading to a net zero effect of spatial activity on group-level sociality (Supporting information **Fig. S5A**).

### (iii) Change in social time predicts individual foraging success (Exp 1)

Change in group-level social time positively correlated with arrival at detected food patches for individuals in highly motivated groups, but not for individuals in lowly motivated groups (Social change*Foraging motivation: χ2 = 4.84, P = 0.03; Supporting information **Table S17**, **Fig. 2**). This relationship between the arrival of individuals at detected food patches and group-level social change did not differ between the sexes (Social change*Sex: χ2 = 2.59, P = 0.11; Supporting information **Table S17**).

**Figure 2.**
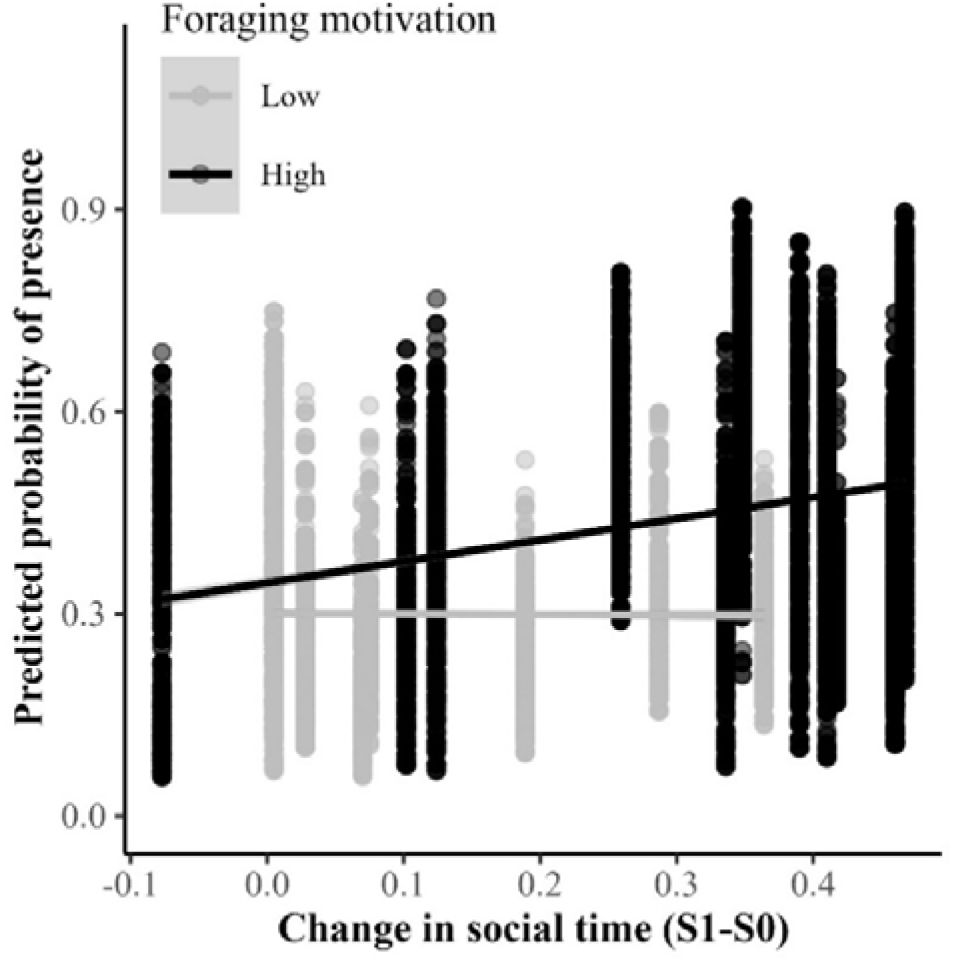
Predicted probability of an individual arriving at a detected food patch in relation to the change in time spent social by an individual’s group following the first series of ephemeral food patch presentations. Predictions were retrieved from the model for individual presence or absence at a discovered food patch. Group-level foraging motivation (proportion of arrived individuals feeding at a patch) is visualised as categorical for illustration purposes (Low: ≤ 0.5, High: > 0.5) but was analysed as a continuous variable. Trend lines were calculated using the function ’geom_smooth()’ from the package ‘ggplot2’.

In both females and males, the proportion of food patches reached positively predicted the number of feeding events (Females: r_s_ = 0.94, N = 69, P < 0.001; Males: r_s_ = 0.90, N = 71, P < 0.001), supporting that patch arrival is indeed a relevant measure of foraging success in this study.

### (iv) Ephemeral food odour patches positively affect social time (Exp 2)

Guppy groups also increased sociality when they could only see and smell patchy ephemeral food resources, but not feed from them (Treatment: χ2 = 13.13, P < 0.001; Supporting information **Table S18**, **Fig. 3**). The average proportion of time spent socially during and after food odour presentations was approximately 1.4 times higher (S1+S2 = **0.44**, 95% CI = 0.38-0.50) compared to control (S1+S2 = **0.31**, 95% CI = 0.26-0.36), indicating that social changes following ephemeral food patch exposure are not merely driven by satiation effects.

**Figure 3.**
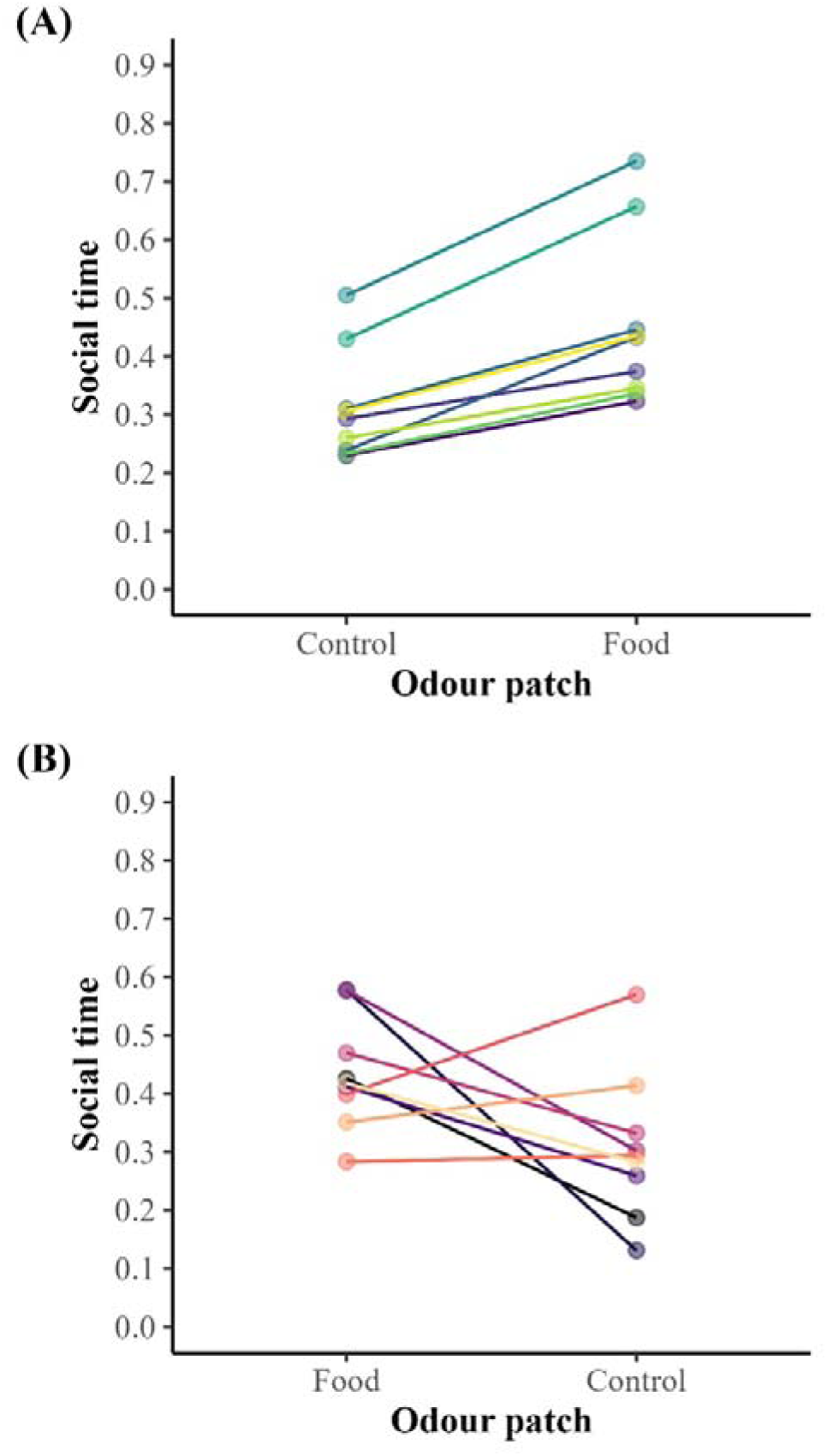
Time spent social by the 18 groups of wild guppies following exposure to two series of ephemeral food odour patch or control presentations. A) Nine groups first received two series of the control treatment, followed by two series of the food odour treatment. B) Nine groups first received two series of the food odour treatment, followed by two series of the control treatment. Social values from the two social sessions (S1 and S2) were combined into one value per group per treatment. Colours depict group ID.

## 4. Discussion

Social flexibility may make animals better prepared to cope with resource fluctuations. Our study provides experimental support for the hypothesis that wild fish can flexibly alter fission-fusion behaviour in response to patchy and ephemeral resource availability, thereby enhancing resource localisation. First, we show that experimentally providing experience with patchy and ephemeral food resources leads to rapid and substantial changes in the time spent social by wild Trinidadian guppies living in resource-poor environments. Our analyses of social and spatial transition probabilities do not support increased spatial activity or local aggregation as the core mechanism underlying this observed increase in sociality. Second, we demonstrate that group-level increase in the overall time spent social positively predicts individual food patch discovery, mediated by group-level foraging motivation. These findings are consistent with an earlier study showing that individual food patch discovery and food consumption of wild guppies increased when the (manipulated) number of guppies in the pool also increased (Snijders et al., 2021). Lastly, experience with patchy and ephemeral food odour stimuli likewise resulted in substantially more time spent social, supporting the hypothesis that the observed social state changes were not merely a consequence of satiation effects. Together, our findings indicate that wild guppies may actively regulate sociality to locate ephemeral food patches better and thus enhance their foraging efficiency in fluctuating environments.

Wild guppy groups doubled their time spent social following the ephemeral presence of novel food patches. This flexible change in social behaviour contrasts with previous environment and density manipulations of guppies in the wild, which did not show such large effects on social time (Krause et al., 2017; Wilson et al., 2015), highlighting the key role of the foraging context in social behaviour. Animals can benefit from the quicker detection of risks and resources in (larger) groups. Indeed, an earlier study showed that more fish in a given pool resulted in quicker detection of ephemeral food patches (Snijders et al., 2021). By spending more time in social proximity, individuals can benefit from such available social information by rapidly detecting social cues about the presence of food resources. Given that wild guppies were shown to join others at a newly discovered food patch within a few seconds (Snijders et al., 2018), one likely social cue is movement initiation or acceleration of a social companion, which subsequently promotes a following response (Franks & Marshall, 2013). In addition, local enhancement likely plays a role (Reader et al., 2003) and local aggregations of feeding conspecifics will likely be more quickly detected by groups than solitary individuals.

Social information use is not always useful but predicted to be particularly valuable for detecting patchy rather than uniformly distributed resources (Oestreich et al 2026), which may explain differences in social response between our study and some earlier (lab) studies (Hoare et al., 2004; Jolles et al., 2018; Schaerf et al., 2017). For example, x-ray tetras (*Pristella maxillaris*) and banded killifish (*Fundulus diaphanus*) reduced encounter frequency and group size, respectively, following exposure to more uniformly presented food cues (Hoare et al., 2004; Schaerf et al., 2017). In addition, variation between species and populations in baseline sociality levels may explain some of the differences with earlier studies. When highly social groups are confronted with patchy resources, they might also do better by initially reducing sociality or becoming more disordered (MacGregor et al., 2020), allowing for the gathering of more independent private information, which can subsequently serve as a source of social information. In contrast, populations with low baseline sociality and minimal aggression, such as our low-predation guppy population, would benefit from increasing sociality to enhance opportunities for acquiring social information.

Recent evidence highlights repeatable yet responsive sociality in animal groups (Jolles et al., 2018; MacGregor and Ioannou, 2022; Schaerf et al., 2017). Our guppy groups showed repeatability in sociality during control presentations, similar to a recent lab study with the same species (Clark et al., 2025). Yet, in our study, between-group variances increased following exposure to ephemeral food patches. We can rule out time or familiarity effects (MacGregor and Ioannou, 2022), as no similar pattern occurred in control treatments. Some groups changed more strongly than others, contrasting with Jolles et al., (2018), who found consistent group structure in sticklebacks (*Gasterosteus aculeatus*) before and after resource depletion. Individual heterogeneity within groups likely played a role, as it can influence group functioning (Farine et al., 2015; Jolles et al., 2020a) and foraging success (Dyer et al., 2009; Hasenjager et al., 2020a). Differences in locomotion and personality types can drive group cohesion, alignment, and leadership (Jolles et al., 2020b, 2017). However, little is known about how group composition affects responsiveness to environmental changes and challenges in the wild (but see Cerritelli et al., 2025). Specific phenotypes may enhance group adaptability, influencing the natural selection of these phenotypes.

Wild animals often need to spend a substantial amount of time locating food resources, constraining the time available for other behaviours, such as sexual pursuits (Borgeaud et al., 2021; Gareta García et al., 2021; Kolluru and Grether, 2005). The increase in sexual behaviour of male guppies following food patch localisation illustrates how increased access to food may influence behavioural time budgets and shows how male guppies, living in resource-limited environments, can enjoy additional reproductive benefits from finding higher-quality food resources (Abrahams, 1993; Kolluru and Grether, 2005). Alternatively, or additionally, the increase in sexual behaviour may have been facilitated by the increase in sociality. Under different circumstances, wild female guppies may generally actively avoid males (Darden and Croft, 2008, but see Snijders et al., 2019). An increase in proximity between males and females thus provides male guppies with more opportunities for sexual pursuits.

To rule out that the observed change in sociality was merely a consequence of satiation-related time-budget changes, rather than an active social strategy, we repeated our experiment using food cues (Hoare et al., 2004; Schaerf et al., 2017), and again observed a substantial increase in time spent socially. Alternatively, a reduction in perceived competition costs following an increase in the temporary (perceived) availability of food could have altered the guppies’ social behaviour. However, the comparably low frequency of aggressive interactions we observed near food and control odour cues suggests relatively high levels of social tolerance (De la Fuente et al., 2022; DeTroy et al., 2021), a population-level characteristic that was also documented in earlier studies (e.g., Snijders et al., 2021). We therefore consider changes in perceived or real competition following our treatments as an unlikely explanation of the substantial changes in sociality we observed.

Group-level responses are not necessarily the sum of individual responses (Bengston and Jandt, 2014), and groups may outperform individuals (Harpaz and Schneidman, 2020; Pitcher et al., 1982; Sasaki et al., 2013). Our experimental design limited our ability to gather robust individual-level data, preventing exploration of whether group-level changes were emergent properties. Future research, with advancements in automated video-tracking, could examine whether changes in group-level sociality predict individual foraging success better than individual-level changes. This may reveal that individuals differ in how much their foraging success improves through social associations versus foraging alone, which could explain individual differences in sociability and its maintenance through selective forces in dynamic resource environments. For example, more social foragers will be more vulnerable to large fluctuations in population density (Harel et al., 2017), while more solitary foragers will generally experience greater variance in foraging, making them more at risk during large fluctuations in available resources (Brown, 1988; Jolles et al., 2017; Oestreich et al., 2026).

In conclusion, through multiple lines of evidence, our study provides new insight into the adaptability of fission-fusion systems under rapidly changing resource conditions in the wild. Specifically, we showed that ephemeral food patch availability generates substantial and profitable changes in an otherwise repeatable group-level social trait. By considering and testing alternative explanations for social change following exposure to temporary food patches, such as carry-over of spatial aggregations, increased spatial activity following food anticipation, sexually motivated behaviours, satiation effects and changes in perceived competition costs, we built a strong case for instrumental use of sociality as a likely explanation. Overall, our findings are consistent with earlier studies in the same population linking increased sociality to increased food patch localisation (Snijders et al., 2021, 2019, 2018). Fish thus appear to be active engineers, rather than passive recipients, of social benefits. Further experimentation is required to reveal the precise social mechanism underlying increased food patch localisation efficiency. Furthermore, revealing the individual and group-level factors driving variation in (adaptive) social responsiveness, the potential mediating effects of individual learning and the relative effects of resource variability on social responsiveness would provide relevant avenues for future research.

## Conflict of Interest

All authors declare no conflict of interest.

## Data availability statement

Supplemental tables and figures can be accessed through the Supporting information file. The data and code are available on the Open Science Framework: https://osf.io/75vtn/overview?view_only=7fb9124575ad447e80ccbb8b6264aa35

## Acknowledgements

We thank Bertrand Jayles for help with the data collection and Eline Weenink for the video analysis. L.S. was funded by a Humboldt Research Fellowship for Postdoctoral Researchers awarded by the Alexander von Humboldt-Stiftung (2019; Ref 3.3 – NLD – 1192631 - HFST-P), supported by the Klaus Tschira Boost Fund (2019; KT09) and a Veni Fellowship awarded by the Netherlands Organisation for Scientific Research (NWO) (2020–2023; VI.Veni.192.018).

## Conflict of Interest

All authors declare no conflict of interest.

## Author contributions

**Lysanne Snijders**: Conceptualization, Data Curation, Formal Analysis, Funding Acquisition, Investigation, Methodology, Project Administration, Resources, Software, Supervision, Validation, Visualization, Writing – original draft, Writing – review & editing; **Stefan Krause**: Conceptualization, Data Curation, Formal Analysis, Investigation, Methodology, Software, Validation, Visualization, Writing – review & editing; **Alan Novaes Tump**: Investigation, Methodology, Writing – review & editing; **Michael Breuker**: Data Curation, Investigation, Software, Writing – review & editing; **Indar W. Ramnarine**: Conceptualization, Project Administration, Resources, Writing – review & editing; **Ralf H.J.M. Kurvers**: Conceptualization, Funding Acquisition, Investigation, Methodology, Project Administration, Supervision, Writing – review & editing; **Jens Krause**: Conceptualization, Funding Acquisition, Investigation, Methodology, Project Administration, Resources, Supervision, Writing – review & editing

## SUPPORTING INFORMATION

### Supporting text

#### Evaluating evidence for local aggregation and effects on social time (Exp 1)

First, to examine whether changes in social time may be strongly biased by the first observed individuals (e.g., following potential temporary carry-over of local aggregation), we tested whether focal observation order (i.e., the order in which individuals were observed during a social session following the food treatment) affected time spent social. For this test, we measured the rank correlation (Kendall’s coefficient) between an individual’s social time and observation order, while performing within-group randomisation of the social times (10,000 randomisation steps) to account for the dependency of within-group social data (**Table S2: *ii*_f**). Individuals were not more or less likely to be social if they were more quickly observed after the first session of food patch presentations (S1: Kendall’s correlation coefficient = 0.03, P = 0.37; **Fig. S2**), although they were slightly more likely to be social when more quickly observed after the second session (S2: Kendall’s correlation coefficient = 0.15, P = 0.01; **Fig. S3**). Overall, we find no indication for strong carry-over of local aggregation and conclude that changes in group-level social time were not overly biased by the first few individuals observed.

Second, we again tested whether carry-over of local aggregation may have influenced our sociality estimates, but now by testing whether a higher social time in Exp 1 was explicitly linked to the location of the last food patch presentation. Specifically, we tested whether the fractions of social contacts in the zone of the last food presentation were higher after the food patch presentations (S1 or S2) than in that same zone before the food patch presentations (S0), using Matched-pairs Wilcoxon tests (**Table S2: *ii*_g**). Social contacts in the zones of the most recently visited food patch were just as likely to take place before as after the temporary food patch became available (**Table S11**), indicating no evidence for carry-over of local aggregation driving the changes in social time.

#### Behavioural interactions at the odour patch (Exp 2)

To evaluate the presence of a potential bias through carry-over effects of food odour patch-related social interactions, we additionally video-scored any social events that took place within two body lengths of the patch: sigmoidal display, inter- and intra-sexual harassment and displacement (departing at the moment another individual arrives). Note, these patch-related social interaction events were not used to quantify the overall social dynamics (see ‘*Focal observations of social dynamics*’). Food odour and control presentations generated a comparable (minor) number of social interactions (sexual and agonistic) at the patch (0.09 interactions per presented patch). When considering only detected (i.e. visited) odour patches, the results were similar (food odour: 0.14 interactions per patch; control: 0.17 interactions per patch) suggesting that any observed increase in sociality following ephemeral food odour patches was not biased by increased social interaction activity near the food odour patch compared to the control.

### Supporting tables

**Table S1.** Definitions of all response variables, explanatory variables and covariates used throughout the analyses. The final column links each variable to one or more of the four main research questions: ***(i)*** repeatability of group sociality; ***(ii)*** effects of ephemeral food on sociality; ***(iii)*** relationship between social change and foraging success; ***(iv)*** effects of perceived ephemeral food availability in the absence of food consumption. P = probability, s = social (within four body lengths), a = alone.

| Variable | Level | Definition | Role | Used in research question(s) |
| --- | --- | --- | --- | --- |
| **Social time** | Group (also sex-within-group in Exp 1) | Proportion of time spent within four body lengths of another guppy, estimated from Markov transition probabilities ($P_{a \rightarrow s}$) / ($P_{s \rightarrow a} + P_{a \rightarrow s}$) | Primary response variable | *(i), (ii), (iv)* |
| **Probability of starting a social contact ($P_{a \rightarrow s}$)** | Group | Probability of transitioning from a solitary state to a non-solitary state at the next sampling point (ten seconds later) | Response (mechanistic analysis) | *(ii)* |
| **Probability of ending a social contact ($P_{s \rightarrow a}$)** | Group | Probability of transitioning from a non-solitary state to solitary state at the next sampling point (ten seconds later) | Response (mechanistic analysis) | *(ii)* |
| **Social session** | Group | Focal observation session of social dynamics before (S0), between (S1) and after (S2) patch presentations | Predictor variable | *(i), (ii)*; S1 and S2 combined and no S0 in *(iv)* |
| **Treatment** | Group | Exp 1: Food patch vs Control patch; Exp 2: Food odour patch vs Control patch | Predictor variable | *(ii) and (iv)*; treatment-specific analyses in *(i) and (iii)* |
| **Treatment sequence** | Group | Food→Control or Control→Food | Predictor variable (control) | *(i)–(iv)* |
| **Pool identity** | Group | Experimental pool (Exp 1: four pools; Exp 2: three pools) | Predictor (control) and nesting variable | *(i)–(iv)* |
| **Group size** | Group | Number of fish in the group (Exp 1: 7–8; Exp 2: 6–8 individuals) | Predictor variable (control) | *(i)–(iv)* |
| **Foraging motivation** | Group | Proportion of fish feeding (calculated as the number of individuals feeding divided by the total number of fish that arrived at each discovered food patch, averaged across all discovered food patches) | Predictor variable (control) | *(i)–(iii)* |
| **Group identity** | Group | Experimental group / batch of guppies receiving the same treatment during the same time in the same pool (Exp 1: 18 groups; Exp 2: 18 groups). | Random effect | *(i)–(iv)* |
| **Sexual behaviour frequency** | Group | Sum of sigmoidal displays and sexual harassment events during focal observations | Response variable | Supplementary analysis for *(ii)* |
| **Observation order** | Individual | Order in which individuals were observed during focal sessions (rank 1 to 8) | Predictor (observation bias test) | Supporting analysis for *(ii)* |
| **Last presentation zone** | Group | Zone of the final food presentation (Exp 1: five zones; Exp 2: nine zones) | Predictor (spatial confound) | Supporting analysis for *(ii)* |
| **Spatial zone transition probability (Spatial activity)** | Group | Probability of transitioning from one spatial zone to another spatial zone at the next sampling point (ten seconds later) | Response and predictor variable (spatial confound) | Supporting analysis for *(ii)* |
| **Arrival at detected food patch** | Individual | Whether an individual reached a food patch that was detected by at least one group member (yes/no) | Response variable | *(iii)* |
| **Social change** | Group | Difference in group social time before (focal session: S0) and in between patch presentations (focal session: S1) | Predictor variable | *(iii)* |
| **Sex** | Individual | Male or female | Predictor variable | *(iii)*; sex-specific analyses in *(ii)* |
| **Body length** | Individual | Standard length (mm), centred within sex | Predictor variable (control) | *(iii)* |
| **Individual identity** | Individual | Marked fish identity (Exp 1: 140 individuals; Exp 2: 137 individuals) | Random effect | *(iii)* |
| **Presentation location** | Patch | Spatial location of patch presentation (Exp 1: five locations/pool; Exp 2: nine locations/pool) | Random effect | *(iii)* |
| **Presentation identity** | Patch | Unique identity of each patch presentation | Random effect | *(iii)* |
| **Feeding events** | Individual | Number of presentations at which an individual consumed food | Response variable (validation) | Supporting analysis for *(iii)* |

**Table S2.**
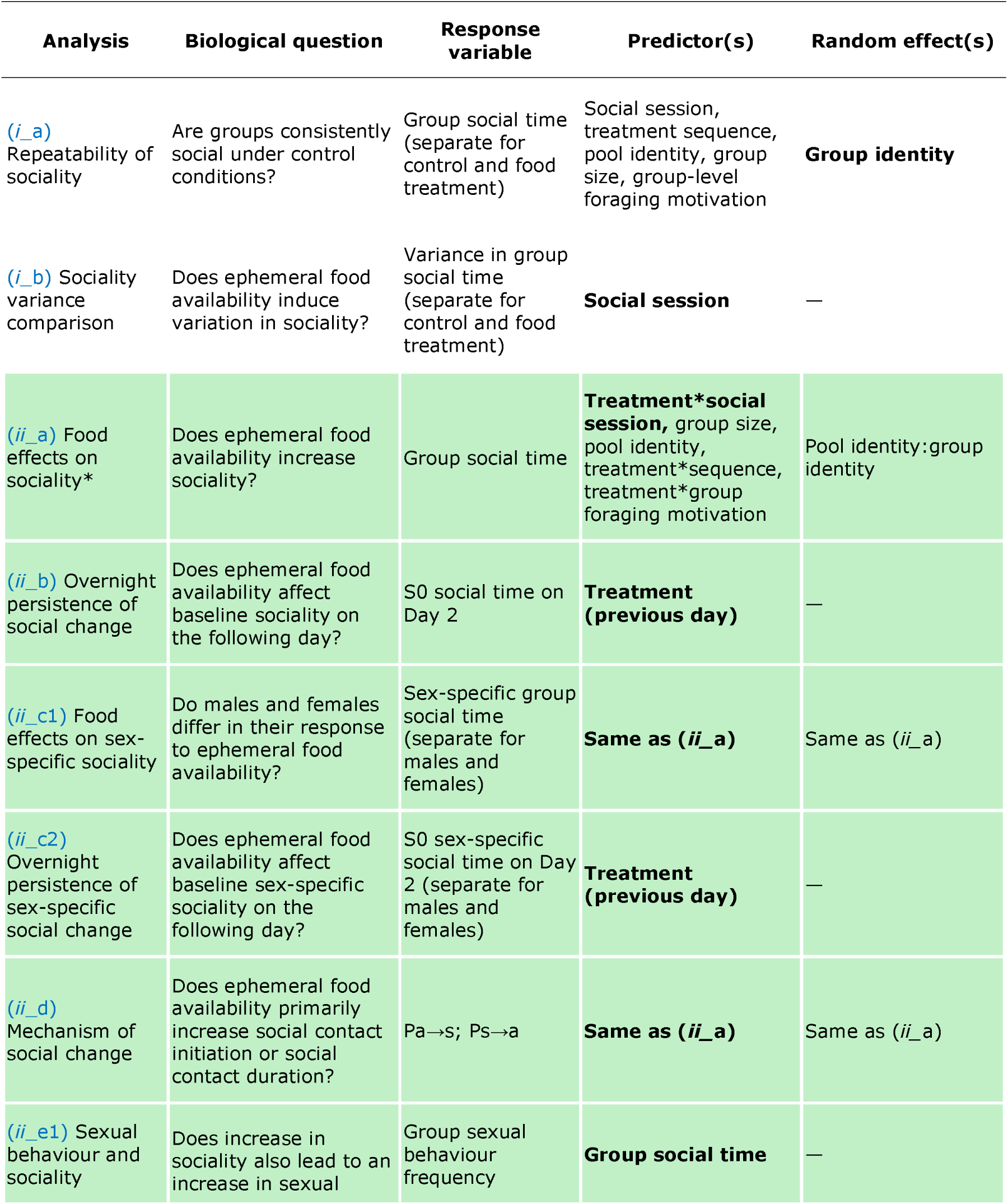
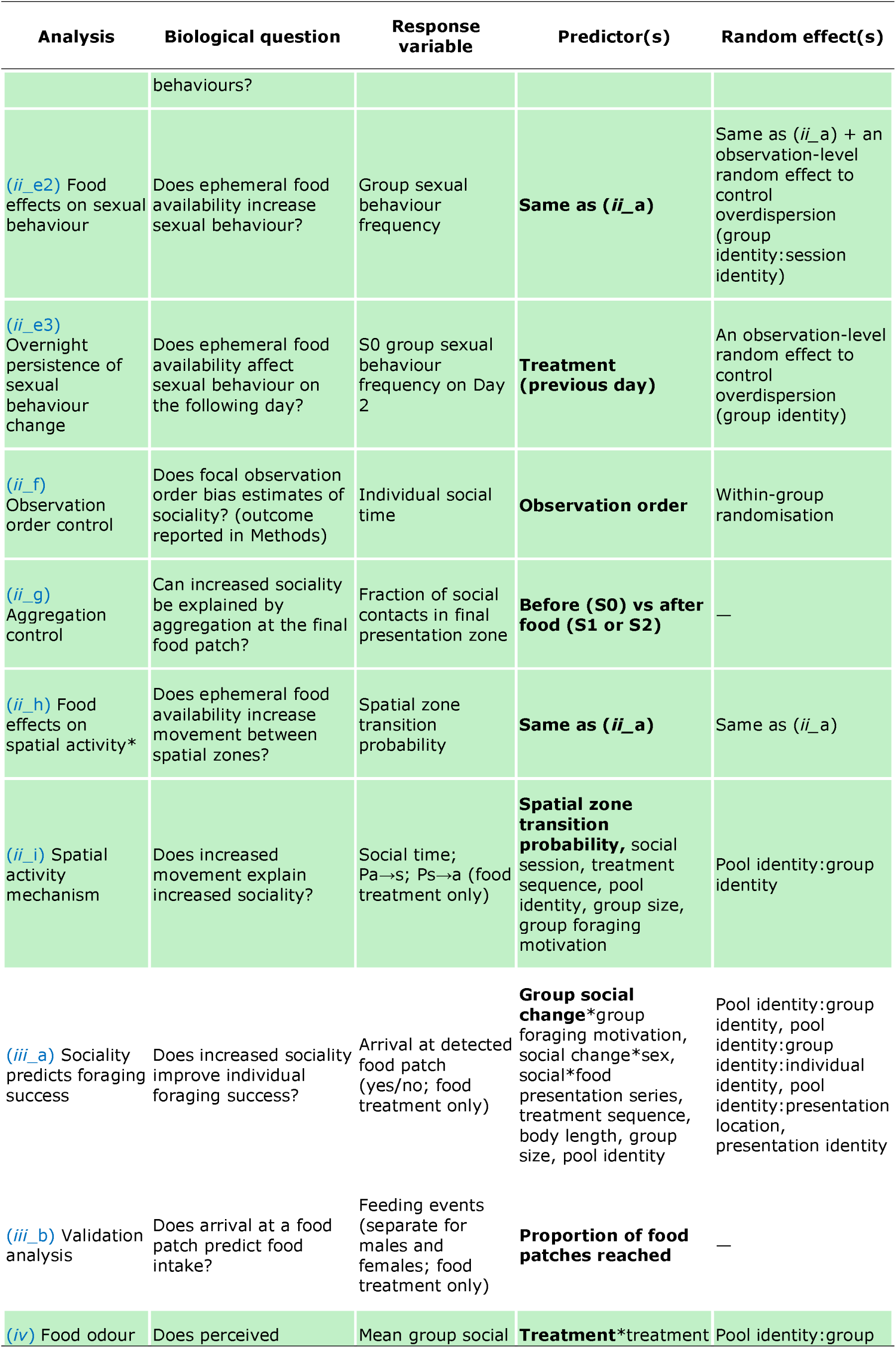
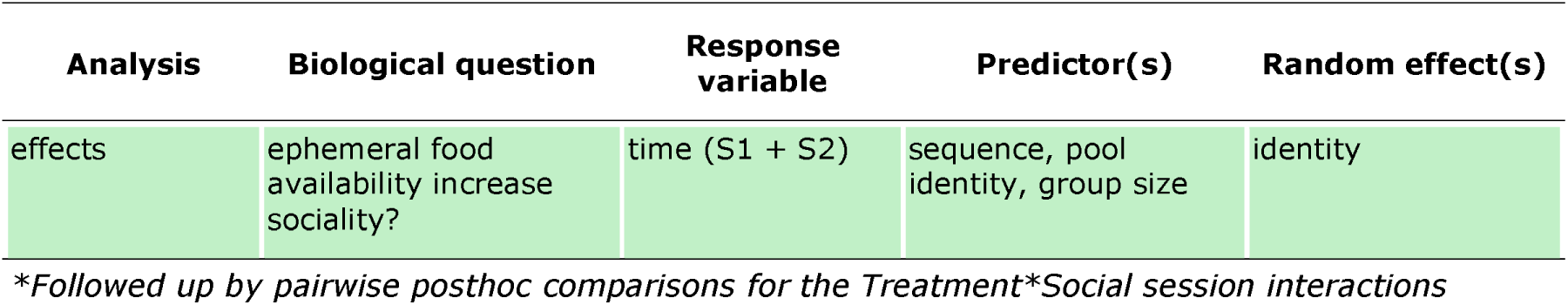
Summary of all statistical analyses, showing the specific biological question addressed, response variables, predictors and random effects. Analysis numbering refers to the four main research question: ***(i)*** repeatability of group sociality; ***(ii)*** effects of ephemeral food (or food odour) on sociality; ***(iii)*** relationship between social change and foraging success; ***(iv)*** effects of perceived ephemeral food availability in the absence of food consumption. P = probability, s = social (within four body lengths), a = alone. Predictors of interest are reported in bold.

**Table S3.** Linear mixed model fixed effects for group-level sociality [Table S2: ii_a]. Continuous variables were centred and scaled. Random effects: Group ID nested in Pool ID (Pool ID:Group ID). Significant estimates are reported in bold.

| Term | Estimate | Std. Error | df | t value | Pr(>\|t\|) |
| --- | --- | --- | --- | --- | --- |
| **(Intercept)** | **0.28** | **0.04** | **32.70** | **6.65** | **0.00** |
| Treatment (Food) | -0.06 | 0.04 | 90.00 | -1.67 | 0.10 |
| Social session (S1) | -0.03 | 0.03 | 90.00 | -1.06 | 0.29 |
| Social session (S2) | -0.02 | 0.03 | 90.00 | -0.53 | 0.60 |
| Sequence (Food-Control) | -0.01 | 0.04 | 33.14 | -0.37 | 0.71 |
| **Foraging motivation** | **-0.05** | **0.02** | **28.95** | **-2.22** | **0.03** |
| Group size (8) | 0.02 | 0.04 | 18.00 | 0.48 | 0.64 |
| Pool (P2) | -0.09 | 0.04 | 18.00 | -2.05 | 0.06 |
| Pool (P3) | 0.03 | 0.03 | 18.00 | 0.82 | 0.42 |
| Pool (P4) | -0.09 | 0.04 | 18.00 | -2.13 | 0.05 |
| **Treatment (Food):Social session (S1)** | **0.27** | **0.04** | **90.00** | **6.32** | **0.00** |
| **Treatment (Food):Social session (S2)** | **0.28** | **0.04** | **90.00** | **6.59** | **0.00** |
| Treatment (Food):Sequence (Food-Control) | 0.03 | 0.04 | 90.00 | 0.77 | 0.44 |
| **Treatment (Food):Foraging motivation** | **0.06** | **0.02** | **90.00** | **3.09** | **0.00** |
| N | 108 |  |  |  |  |
| AIC | -169.43 |  |  |  |  |
| BIC | -126.51 |  |  |  |  |
| Log likelihood | 100.71 |  |  |  |  |

**Table S4.** Post-hoc contrasts* for the significant interaction between treatment and social session in the linear mixed model for group-level sociality [Table S2: ii_a]. Significant estimates are reported in bold.

| Social session | Treatment | Contrast | Estimate | Std. Error | df | t-ratio | P |
| --- | --- | --- | --- | --- | --- | --- | --- |
| S0 |  | Food-Control | -0.04 | 0.03 | 96.4 | -1.44 | 0.58 |
| **S1** |  | **Food-Control** | **0.22** | **0.03** | **96.4** | **7.17** | **0.00** |
| **S2** |  | **Food-Control** | **0.23** | **0.03** | **96.4** | **7.54** | **0.00** |
|  | Control | S1-S0 | -0.03 | 0.03 | **96.4** | -1.02 | 0.85 |
|  | Control | S2-S1 | 0.02 | 0.03 | **96.4** | 0.51 | 0.99 |
|  | **Food** | **S1-S0** | **0.24** | **0.03** | **96.4** | **7.58** | **0.00** |
|  | Food | S2-S1 | 0.03 | 0.03 | **96.4** | 0.88 | 0.91 |

\* Results are averaged over some or all of the levels of: Treatment sequence, Group size, Pool identity. Degrees-of-freedom method: kenward-roger P value adjustment: mvt method for 7 tests.

**Table S5.** Linear mixed model fixed effects for female group-level sociality [Table S2: ii_c]. Continuous variables were centred and scaled. Random effects: Group ID nested in Pool ID (Pool ID:Group ID). Significant estimates are reported in bold.

| Term | Estimate | Std. Error | df | t value | Pr(>\|t\|) |
| --- | --- | --- | --- | --- | --- |
| **(Intercept)** | **0.27** | **0.05** | **40.55** | **5.72** | **0.00** |
| Treatment (Food) | -0.07 | 0.05 | 90.00 | -1.51 | 0.14 |
| Social session (S1) | -0.02 | 0.04 | 90.00 | -0.48 | 0.63 |
| Social session (S2) | 0.02 | 0.04 | 90.00 | 0.57 | 0.57 |
| Foraging motivation | -0.01 | 0.02 | 34.80 | -0.57 | 0.57 |
| Treatment sequence (Food-Control) | 0.01 | 0.04 | 41.21 | 0.23 | 0.82 |
| Group size (8) | 0.01 | 0.04 | 18.00 | 0.28 | 0.78 |
| Pool (P2) | -0.01 | 0.05 | 18.00 | -0.22 | 0.83 |
| Pool (P3) | 0.02 | 0.03 | 18.00 | 0.46 | 0.65 |
| Pool (P4) | -0.08 | 0.04 | 18.00 | -1.87 | 0.08 |
| **Treatment (Food):Social session (S1)** | **0.24** | **0.06** | **90.00** | **4.42** | **0.00** |
| **Treatment (Food):Social session (S2)** | **0.26** | **0.06** | **90.00** | **4.68** | **0.00** |
| Treatment (Food):Foraging motivation | 0.05 | 0.03 | 90.00 | 1.85 | 0.07 |
| Treatment (Food):Treatment sequence (Food-Control) | 0.06 | 0.05 | 90.00 | 1.13 | 0.26 |
| N | 108 |  |  |  |  |
| AIC | -119.76 |  |  |  |  |
| BIC | -76.85 |  |  |  |  |
| Log likelihood | 75.88 |  |  |  |  |

**Table S6.**
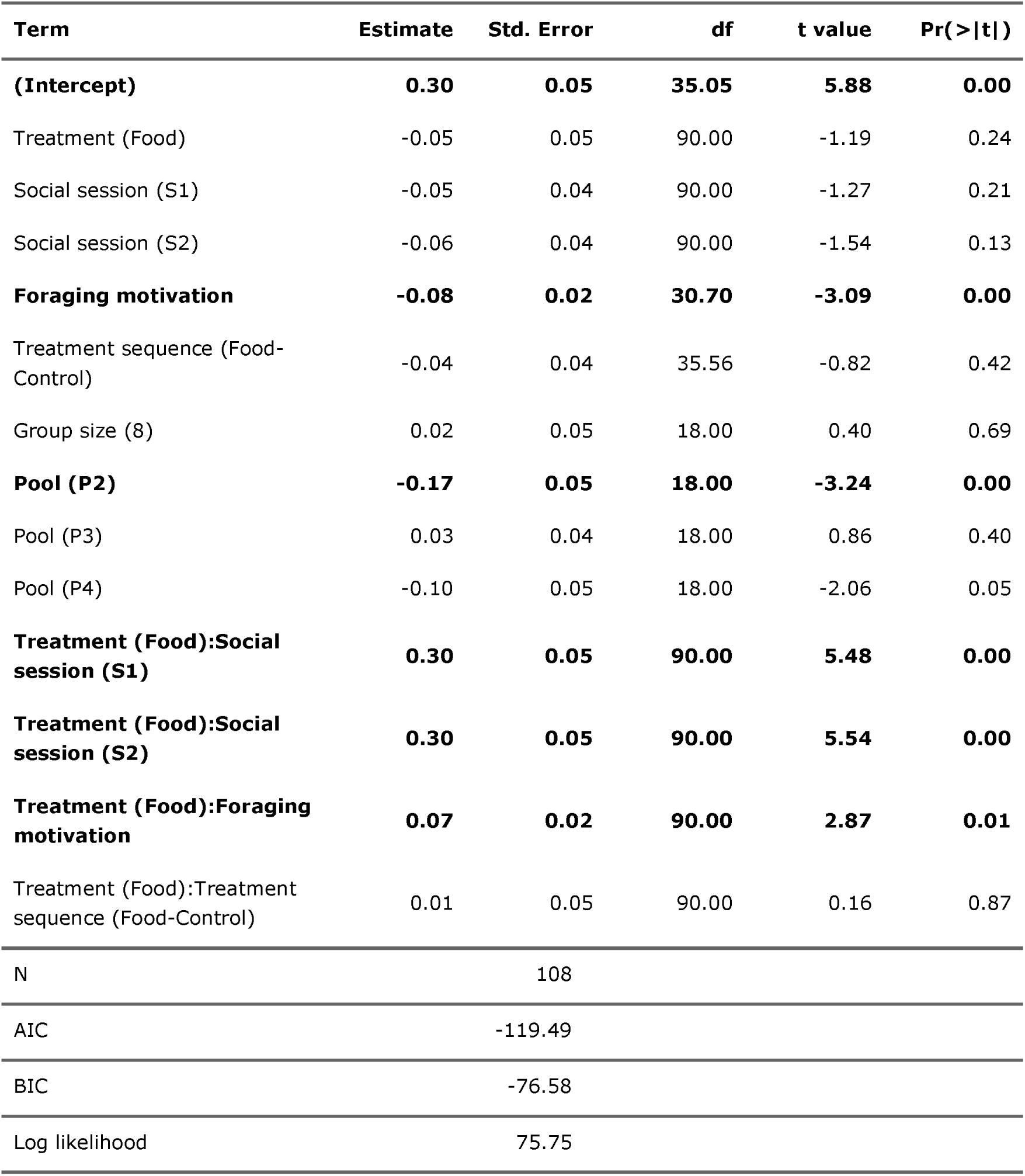
Linear mixed model fixed effects for male group-level sociality [Table S2: ii_c]. Continuous variables were centred and scaled. Random effects: Group ID nested in Pool ID (Pool ID:Group ID). Significant estimates are reported in bold.

| Term | Estimate | Std. Error | df | t value | Pr(>\|t\|) |
| --- | --- | --- | --- | --- | --- |
| **(Intercept)** | **0.30** | **0.05** | **35.05** | **5.88** | **0.00** |
| Treatment (Food) | -0.05 | 0.05 | 90.00 | -1.19 | 0.24 |
| Social session (S1) | -0.05 | 0.04 | 90.00 | -1.27 | 0.21 |
| Social session (S2) | -0.06 | 0.04 | 90.00 | -1.54 | 0.13 |
| **Foraging motivation** | **-0.08** | **0.02** | **30.70** | **-3.09** | **0.00** |
| Treatment sequence (Food-Control) | -0.04 | 0.04 | 35.56 | -0.82 | 0.42 |
| Group size (8) | 0.02 | 0.05 | 18.00 | 0.40 | 0.69 |
| **Pool (P2)** | **-0.17** | **0.05** | **18.00** | **-3.24** | **0.00** |
| Pool (P3) | 0.03 | 0.04 | 18.00 | 0.86 | 0.40 |
| Pool (P4) | -0.10 | 0.05 | 18.00 | -2.06 | 0.05 |
| **Treatment (Food):Social session (S1)** | **0.30** | **0.05** | **90.00** | **5.48** | **0.00** |
| **Treatment (Food):Social session (S2)** | **0.30** | **0.05** | **90.00** | **5.54** | **0.00** |
| **Treatment (Food):Foraging motivation** | **0.07** | **0.02** | **90.00** | **2.87** | **0.01** |
| Treatment (Food):Treatment sequence (Food-Control) | 0.01 | 0.05 | 90.00 | 0.16 | 0.87 |
| N | 108 |  |  |  |  |
| AIC | -119.49 |  |  |  |  |
| BIC | -76.58 |  |  |  |  |
| Log likelihood | 75.75 |  |  |  |  |

**Table S7.** Linear mixed model fixed effects for the group-level transition probability from alone to social (Pa→s) [Table S2: ii_d]. Continuous variables were centred and scaled. Random effects: Group ID nested in Pool ID (Pool ID:Group ID). Significant estimates are reported in bold.

| Term | Estimate | Std. Error | df | t value | Pr(>\|t\|) |
| --- | --- | --- | --- | --- | --- |
| **(Intercept)** | **0.14** | **0.02** | **47.44** | **6.82** | **0.00** |
| Treatment (Food) | -0.02 | 0.02 | 90.00 | -0.69 | 0.50 |
| Social session (S1) | -0.02 | 0.02 | 90.00 | -1.05 | 0.30 |
| Social session (S2) | 0.00 | 0.02 | 90.00 | -0.17 | 0.87 |
| Foraging motivation | -0.02 | 0.01 | 40.03 | -1.57 | 0.12 |
| Treatment sequence (Food-Control) | 0.00 | 0.02 | 48.28 | 0.09 | 0.93 |
| Group size (8) | 0.00 | 0.02 | 18.00 | 0.24 | 0.81 |
| Pool (P2) | -0.01 | 0.02 | 18.00 | -0.57 | 0.58 |
| Pool (P3) | 0.00 | 0.01 | 18.00 | -0.34 | 0.74 |
| Pool (P4) | 0.00 | 0.02 | 18.00 | -0.15 | 0.89 |
| **Treatment (Food):Social session (S1)** | **0.14** | **0.03** | **90.00** | **5.28** | **0.00** |
| **Treatment (Food):Social session (S2)** | **0.09** | **0.03** | **90.00** | **3.42** | **0.00** |
| **Treatment (Food):Foraging motivation** | **0.03** | **0.01** | **90.00** | **2.14** | **0.04** |
| Treatment (Food):Treatment sequence (Food-Control) | 0.00 | 0.02 | 90.00 | 0.11 | 0.92 |
| N | 108 |  |  |  |  |
| AIC | -282.93 |  |  |  |  |
| BIC | -240.02 |  |  |  |  |
| Log likelihood | 157.47 |  |  |  |  |

**Table S8.** Linear mixed model fixed effects for the group-level transition probability from social to alone (Ps→a)) [Table S2: ii_d]. Continuous variables were centred and scaled. Random effects: Group ID nested in Pool ID (Pool ID:Group ID). Significant estimates are reported in bold.

| Term | Estimate | Std. Error | df | t value | Pr(>\|t\|) |
| --- | --- | --- | --- | --- | --- |
| **(Intercept)** | **0.38** | **0.06** | **31.24** | **6.75** | **0.00** |
| Treatment (Food) | 0.07 | 0.05 | 90.00 | 1.52 | 0.13 |
| Social session (S1) | -0.01 | 0.04 | 90.00 | -0.16 | 0.88 |
| Social session (S2) | 0.01 | 0.04 | 90.00 | 0.27 | 0.79 |
| Foraging motivation | 0.04 | 0.03 | 27.87 | 1.36 | 0.19 |
| Treatment sequence (Food-Control) | 0.03 | 0.05 | 31.63 | 0.59 | 0.56 |
| Group size (8) | -0.03 | 0.06 | 18.00 | -0.52 | 0.61 |
| **Pool (P2)** | **0.15** | **0.06** | **18.00** | **2.52** | **0.02** |
| Pool (P3) | -0.01 | 0.04 | 18.00 | -0.14 | 0.89 |
| **Pool (P4)** | **0.19** | **0.06** | **18.00** | **3.40** | **0.00** |
| **Treatment (Food):Social session (S1)** | **-0.18** | **0.06** | **90.00** | **-3.23** | **0.00** |
| **Treatment (Food):Social session (S2)** | **-0.24** | **0.06** | **90.00** | **-4.38** | **0.00** |
| Treatment (Food):Foraging motivation | -0.02 | 0.03 | 90.00 | -0.97 | 0.34 |
| Treatment (Food):Treatment sequence (Food-Control) | 0.00 | 0.05 | 90.00 | 0.05 | 0.96 |
| N | 108 |  |  |  |  |
| AIC | -109.66 |  |  |  |  |
| BIC | -66.74 |  |  |  |  |
| Log likelihood | 70.83 |  |  |  |  |

**Table S9.** Generalized linear mixed model fixed effects group-level sexual behaviour (display and harassment) [Table S2: ii_e2]. Continuous variables were centred and scaled. Random effects: Group ID nested in Pool ID (Pool ID:Group ID) and an observation-level random effect to treat overdispersion (Group ID:Session ID). Family = Poisson. Significant effects are reported in bold.

| Term | Estimate | SE | Statistic | p |
| --- | --- | --- | --- | --- |
| **(Intercept)** | **2.01** | **0.27** | **7.33** | **0.00** |
| Treatment (Food) | -0.24 | 0.27 | -0.89 | 0.37 |
| Social session (S1) | -0.09 | 0.22 | -0.40 | 0.69 |
| Social session (S2) | 0.03 | 0.22 | 0.12 | 0.91 |
| Foraging motivation | -0.08 | 0.13 | -0.60 | 0.55 |
| Treatment sequence (Food-Control) | 0.24 | 0.24 | 1.03 | 0.31 |
| Group size (8) | 0.19 | 0.25 | 0.74 | 0.46 |
| Pool (P2) | -0.07 | 0.26 | -0.26 | 0.80 |
| Pool (P3) | -0.30 | 0.18 | -1.64 | 0.10 |
| **Pool (P4)** | **-1.16** | **0.27** | **-4.37** | **0.00** |
| **Treatment (Food):Social session (S1)** | **1.10** | **0.31** | **3.58** | **0.00** |
| **Treatment (Food):Social session (S2)** | **1.04** | **0.31** | **3.41** | **0.00** |
| Treatment (Food):Foraging motivation | 0.18 | 0.14 | 1.28 | 0.20 |
| Treatment (Food):Treatment sequence (Food-Control) | -0.06 | 0.27 | -0.21 | 0.84 |
| N | 54 |  |  |  |
| AIC | -47.62 |  |  |  |
| BIC | -23.75 |  |  |  |
| Log likelihood | 35.81 |  |  |  |

**Table S10.** Post-hoc contrasts* for the significant interaction between treatment and social session in the linear mixed model for group-level sexual behaviour [Table S2: ii_e]. Significant estimates are reported in bold.

| Social session | Treatment | Contrast | Estimate | Std. Error | df | z-ratio | P |
| --- | --- | --- | --- | --- | --- | --- | --- |
| S0 |  | Food-Control | -0.27 | 0.22 | Inf | -1.19 | 0.75 |
| **S1** |  | **Food-Control** | **0.84** | **0.21** | Inf | **3.95** | **<0.001** |
| **S2** |  | **Food-Control** | **0.78** | **0.21** | Inf | **3.72** | **0.001** |
|  | Control | S1-S0 | -0.08 | 0.22 | Inf | -0.40 | >0.99 |
|  | Control | S2-S1 | 0.11 | 0.22 | Inf | 0.52 | 0.99 |
|  | **Food** | **S1-S0** | **1.01** | **0.22** | Inf | **4.72** | **<0.001** |
|  | Food | S2-S1 | 0.05 | 0.20 | Inf | 0.27 | >0.99 |

\* Results are averaged over some or all of the levels of: Treatment sequence, Group size, Pool identity. Results are given on the log (not the response) scale. P value adjustment: mvt method for 7 tests.

**Table S11.**
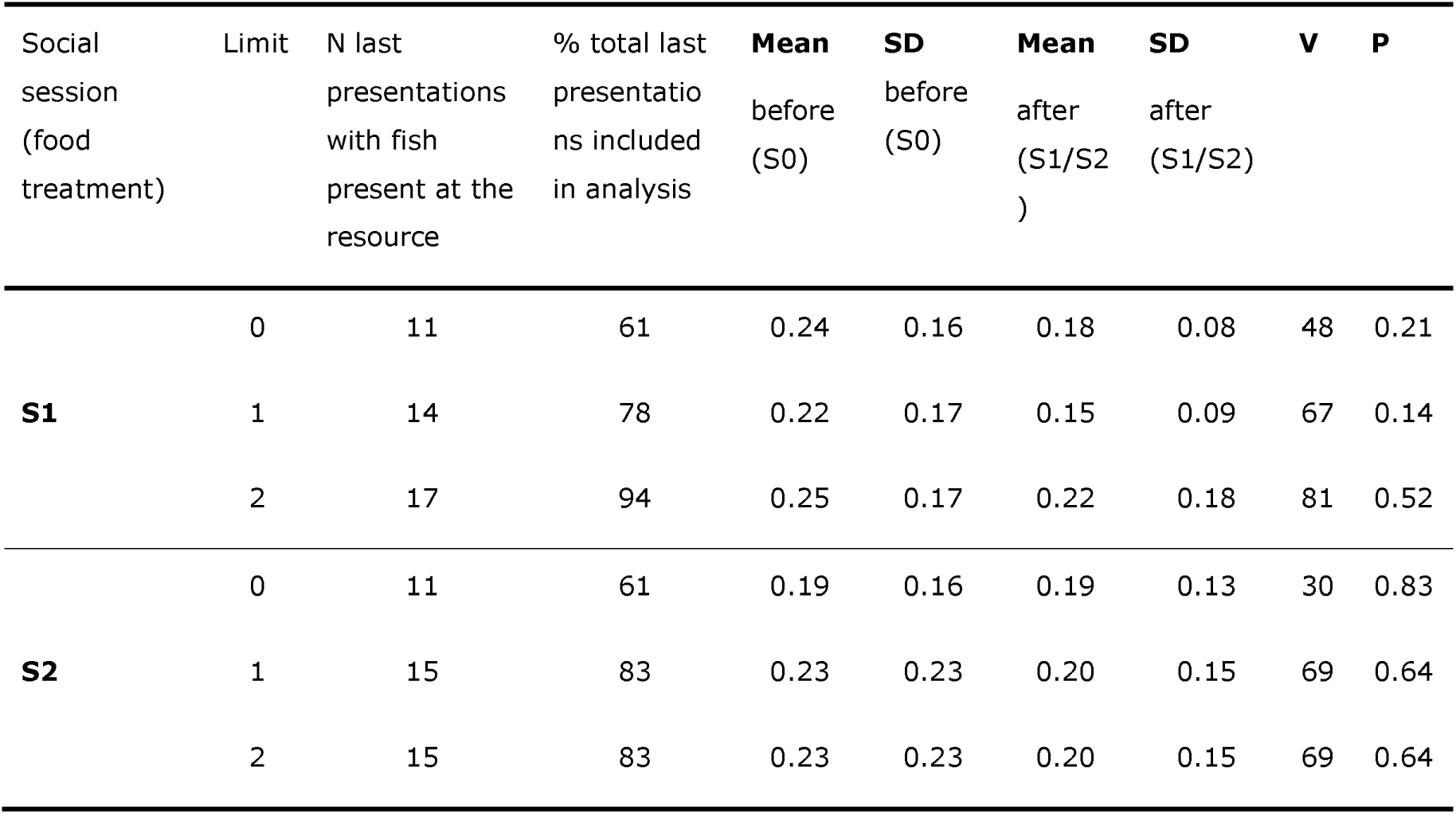
Wilcoxon tests for social carry-over effects of local aggregation at the spatial zone of the last food presentation [Table S2: ii_g]. Relative presence at the zone which fish last visited during the food presentations was compared between before (S0) and after the presentations (S1 or S2). A limit of 0 means only the final presentation in a session was considered and only included if fish were present at the resource. A limit of 1 (or 2) means that, in case no fish were present at the resource during the final presentation, also the second (or third) to last presentation was considered, and included if fish were present there.

| Social session (food treatment) | Limit | N last presentations with fish present at the resource | % total last presentations included in analysis | Mean before (S0) | SD before (S0) | Mean after (S1/S2) | SD after (S1/S2) | V | P |
| --- | --- | --- | --- | --- | --- | --- | --- | --- | --- |
| **S1** | 0 | 11 | 61 | 0.24 | 0.16 | 0.18 | 0.08 | 48 | 0.21 |
|  | 1 | 14 | 78 | 0.22 | 0.17 | 0.15 | 0.09 | 67 | 0.14 |
|  | 2 | 17 | 94 | 0.25 | 0.17 | 0.22 | 0.18 | 81 | 0.52 |
| **S2** | 0 | 11 | 61 | 0.19 | 0.16 | 0.19 | 0.13 | 30 | 0.83 |
|  | 1 | 15 | 83 | 0.23 | 0.23 | 0.20 | 0.15 | 69 | 0.64 |
|  | 2 | 15 | 83 | 0.23 | 0.23 | 0.20 | 0.15 | 69 | 0.64 |

**Table S12.** Linear mixed model fixed effects for the group-level spatial zone transition probability [Table S2: ii_h]. Continuous variables were centred and scaled. Random effects: Group ID nested in Pool ID (Pool ID:Group ID). Significant estimates are reported in bold.

| Term | Estimate | Std. Error | df | t value | Pr(>\|t\|) |
| --- | --- | --- | --- | --- | --- |
| **(Intercept)** | **0.15** | **0.02** | **39.12** | **6.43** | **0.00** |
| Treatment (Food) | 0.00 | 0.02 | 90.00 | -0.16 | 0.87 |
| Social session (S1) | -0.01 | 0.02 | 90.00 | -0.71 | 0.48 |
| Social session (S2) | 0.00 | 0.02 | 90.00 | 0.14 | 0.89 |
| Foraging motivation | -0.01 | 0.01 | 33.73 | -0.54 | 0.59 |
| Treatment sequence (Food-Control) | 0.02 | 0.02 | 39.74 | 1.11 | 0.27 |
| Group size (8) | 0.02 | 0.02 | 18.00 | 0.78 | 0.45 |
| Pool (P2) | 0.00 | 0.02 | 18.00 | -0.15 | 0.88 |
| Pool (P3) | -0.03 | 0.02 | 18.00 | -2.10 | 0.05 |
| Pool (P4) | 0.03 | 0.02 | 18.00 | 1.31 | 0.21 |
| **Treatment (Food):Social session (S1)** | **0.07** | **0.03** | **90.00** | **2.76** | **0.01** |
| Treatment (Food):Social session (S2) | 0.05 | 0.03 | 90.00 | 1.88 | 0.06 |
| **Treatment (Food):Foraging motivation** | **0.03** | **0.01** | **90.00** | **2.75** | **0.01** |
| Treatment (Food):Treatment sequence (Food-Control) | 0.02 | 0.02 | 90.00 | 0.86 | 0.39 |
| N | 108 |  |  |  |  |
| AIC | -280.21 |  |  |  |  |
| BIC | -237.30 |  |  |  |  |
| Log likelihood | 156.11 |  |  |  |  |

**Table S13.** Post-hoc contrasts* for the significant interaction between treatment and social session in the linear mixed model for group-level spatial zone transition probability [Table S2: ii_i]. Significant effects are reported in bold.

| Social session | Treatment | Contrast | Estimate | SE | DF | t-ratio | P |
| --- | --- | --- | --- | --- | --- | --- | --- |
| S0 |  | Food-Control | 0.01 | 0.02 | 97.6 | 0.35 | >0.99 |
| **S1** |  | **Food-Control** | **0.08** | **0.02** | **97.6** | **4.09** | **0.007** |
| **S2** |  | **Food-Control** | **0.06** | **0.02** | **97.6** | **2.89** | **0.03** |
|  | Control | S1-S0 | -0.01 | 0.02 | 97.6 | -0.68 | 0.97 |
|  | Control | S2-S1 | 0.02 | 0.02 | 97.6 | 0.81 | 0.93 |
|  | **Food** | **S1-S0** | **0.06** | **0.02** | **97.6** | **3.06** | **0.02** |
|  | Food | S2-S1 | -0.01 | 0.02 | 97.6 | -0.38 | >0.99 |

\* Results are averaged over some or all of the levels of: Treatment sequence, Group size, Pool identity. Degrees-of-freedom method: kenward-roger P value adjustment: mvt method for 7 tests.

**Table S14.** Linear mixed model summary analysing the effect of group-level spatial zone transition probability (spatial activity) on group-level sociality during food treatment [Table S2: ii_i]. Continuous variables were centred and scaled. Random effects: Group ID nested in Pool ID (Pool ID:Group ID). Significant estimates are reported in bold.

| Term | Estimate | Std. Error | df | t value | Pr(>\|t\|) |
| --- | --- | --- | --- | --- | --- |
| **(Intercept)** | **0.23** | **0.06** | **35.35** | **3.65** | **0.00** |
| Spatial zone transition probability | 0.08 | 0.25 | 53.24 | 0.31 | 0.76 |
| Foraging motivation | 0.02 | 0.02 | 18.41 | 0.78 | 0.44 |
| Group size (8) | -0.04 | 0.05 | 16.70 | -0.70 | 0.49 |
| **Social session (S1)** | **0.23** | **0.04** | **38.77** | **5.79** | **0.00** |
| **Social session (S2)** | **0.26** | **0.04** | **37.88** | **6.60** | **0.00** |
| Treatment sequence (Food-Control) | 0.03 | 0.04 | 18.30 | 0.64 | 0.53 |
| Pool (P2) | -0.08 | 0.06 | 16.69 | -1.47 | 0.16 |
| Pool (P3) | 0.07 | 0.04 | 18.12 | 1.77 | 0.09 |
| **Pool (P4)** | **-0.12** | **0.05** | **16.85** | **-2.35** | **0.03** |
| N | 54.00 |  |  |  |  |
| AIC | -59.12 |  |  |  |  |
| BIC | -35.26 |  |  |  |  |
| Log likelihood | 41.56 |  |  |  |  |

**Table S15.** Linear mixed model summary analysing the effect of group-level spatial zone transition probability (spatial activity) on group-level probability to transition from alone to social (Pa→s) during food treatment [Table S2: ii_i]. Continuous variables were centred and scaled. Random effects: Group ID nested in Pool ID (Pool ID:Group ID). Significant estimates are reported in bold.

| Term | Estimate | Std. Error | df | t value | Pr(>\|t\|) |
| --- | --- | --- | --- | --- | --- |
| (Intercept) | 0.06 | 0.03 | 54 | 1.70 | 0.10 |
| **Spatial zone transition probability** | **0.58** | **0.13** | **54** | **4.31** | **0.00** |
| Foraging motivation | -0.01 | 0.01 | 54 | -0.56 | 0.58 |
| Group size (8) | -0.02 | 0.03 | 54 | -0.58 | 0.56 |
| **Social session (S1)** | **0.09** | **0.02** | **54** | **4.03** | **0.00** |
| **Social session (S2)** | **0.06** | **0.02** | **54** | **2.74** | **0.01** |
| Treatment sequence (Food-Control) | -0.02 | 0.02 | 54 | -0.88 | 0.38 |
| Pool (P2) | -0.02 | 0.03 | 54 | -0.84 | 0.41 |
| Pool (P3) | 0.02 | 0.02 | 54 | 0.71 | 0.48 |
| Pool (P4) | -0.03 | 0.03 | 54 | -1.18 | 0.24 |
| N | 54 |  |  |  |  |
| AIC | -127.61 |  |  |  |  |
| BIC | -103.74 |  |  |  |  |
| Log likelihood | 75.80 |  |  |  |  |

**Table S16.** Linear mixed model summary analysing the effect of group-level spatial zone transition probability (spatial activity) on group-level probability to transition from social to alone (Ps→a) during food treatment [Table S2: ii_i]. Continuous variables were centred and scaled. Random effects: Group ID nested in Pool ID (Pool ID:Group ID). Significant estimates are reported in bold.

| Term | Estimate | Std. Error | df | t value | Pr(>\|t\|) |
| --- | --- | --- | --- | --- | --- |
| **(Intercept)** | **0.36** | **0.07** | **54** | **5.14** | **0.00** |
| **Spatial zone transition probability** | **0.61** | **0.28** | **54** | **2.15** | **0.04** |
| Foraging motivation | -0.02 | 0.03 | 54 | -0.89 | 0.38 |
| Group size (8) | 0.02 | 0.06 | 54 | 0.42 | 0.68 |
| **Social session (S1)** | **-0.22** | **0.04** | **54** | **-4.93** | **0.00** |
| **Social session (S2)** | **-0.26** | **0.04** | **54** | **-5.98** | **0.00** |
| Treatment sequence (Food-Control) | -0.02 | 0.05 | 54 | -0.37 | 0.71 |
| Pool (P2) | 0.08 | 0.06 | 54 | 1.25 | 0.22 |
| Pool (P3) | -0.05 | 0.04 | 54 | -1.04 | 0.30 |
| **Pool (P4)** | **0.18** | **0.06** | **54** | **3.11** | **0.00** |
| N | 54 |  |  |  |  |
| AIC | -47.62 |  |  |  |  |
| BIC | -23.75 |  |  |  |  |
| Log likelihood | 35.81 |  |  |  |  |

**Table S17.** Generalized linear mixed model fixed effects for individual arrival at a detected food patch [Table S2: iii_a]. Continuous variables were centred and scaled. Body length was centred on sex. Random effects: Group ID nested in Pool (Pool:Group ID), Individual ID nested in Group ID (Pool:Group ID:Individual ID), Presentation location nested in Pool (Pool:Presentation location) and Presentation ID. Family = Binomial. Significant effects are reported in bold.

| Term | Estimate | SE | Statistic | p |
| --- | --- | --- | --- | --- |
| (Intercept) | 0.00 | 0.27 | 0.00 | 1.00 |
| **Social change** | **0.27** | **0.12** | **2.37** | **0.02** |
| **Foraging motivation** | **0.41** | **0.10** | **4.02** | **0.00** |
| Sex (Male) | -0.09 | 0.11 | -0.78 | 0.43 |
| Presentation series (F2) | 0.00 | 0.06 | -0.03 | 0.98 |
| Treatment sequence (Food-Control) | -0.18 | 0.21 | -0.83 | 0.41 |
| **Body length** | **0.13** | **0.06** | **2.24** | **0.02** |
| Group size (8) | -0.43 | 0.22 | -1.93 | 0.05 |
| Pool (P2) | -0.49 | 0.37 | -1.33 | 0.18 |
| Pool (P3) | -0.57 | 0.31 | -1.86 | 0.06 |
| Pool (P4) | -0.23 | 0.34 | -0.68 | 0.50 |
| **Social change:Foraging motivation** | **0.28** | **0.13** | **2.20** | **0.03** |
| Social change:Sex (Male) | -0.18 | 0.11 | -1.61 | 0.11 |
| Social change:Presentation series (F2) | -0.05 | 0.06 | -0.91 | 0.36 |
| N | 9,663 |  |  |  |
| AIC | 11,812.39 |  |  |  |
| BIC | 11,941.56 |  |  |  |
| Log likelihood | -5,888.19 |  |  |  |

**Table S18.** Linear mixed model fixed effects of food odour patch treatment on group-level sociality [Table S2: iv]. Social time was averaged across S1 and S2. Random effects: Group ID.

| Term | Estimate | Std. Error | df | t value | Pr(>\|t\|) |
| --- | --- | --- | --- | --- | --- |
| **(Intercept)** | **0.29** | **0.07** | **36** | **4.36** | **0.00** |
| **Treatment (Food)** | **0.14** | **0.04** | **36** | **3.62** | **0.00** |
| Treatment sequence (Food-Control) | 0.00 | 0.04 | 36 | 0.03 | 0.98 |
| Pool (P2) | 0.01 | 0.03 | 36 | 0.39 | 0.70 |
| **Pool (P3)** | **0.16** | **0.04** | **36** | **3.62** | **0.00** |
| Group size (7) | -0.08 | 0.06 | 36 | -1.39 | 0.17 |
| Group size (8) | -0.03 | 0.06 | 36 | -0.57 | 0.58 |
| Treatment (Food):Treatment sequence (Food-Control) | -0.01 | 0.06 | 36 | -0.27 | 0.79 |
| N | 36 |  |  |  |  |
| AIC | -56.91 |  |  |  |  |
| BIC | -41.08 |  |  |  |  |
| Log likelihood | 38.46 |  |  |  |  |

### Supporting figures

**Figure S1.**
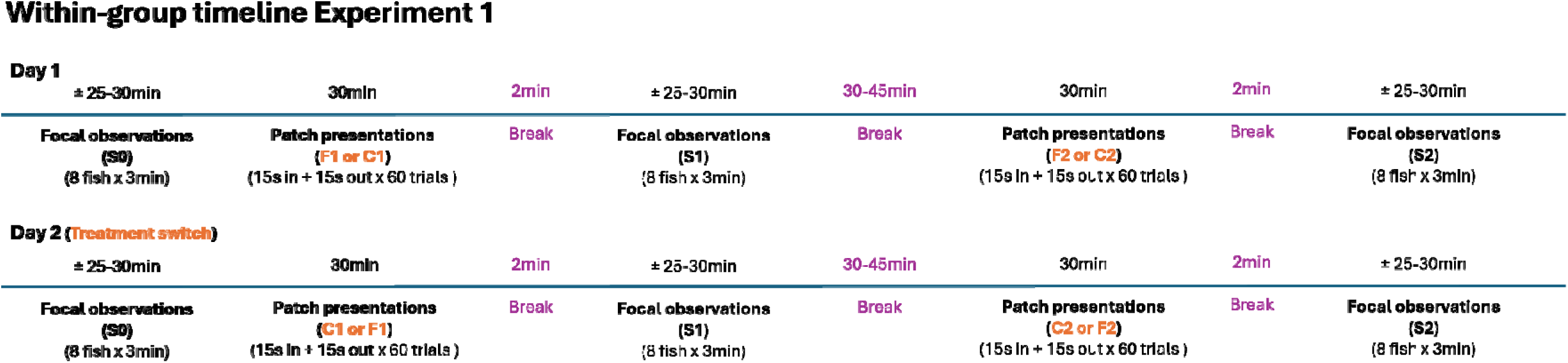
Detailed within group timeline for experiment 1 and experiment 2. In both experiments, groups were exposed to two sequences of food (or odour) treatment (F1 and F2) and two sequences of control treatment (C1 and C2). In experiment 1, groups experienced one treatment on the first day and switched to the other treatment on the second day. In experiment 2, groups experienced one treatment during the first part of the day and switched to the second treatment during the second part of the same day. Focal observations of social behaviour were conducted before (S0), in between (S1) and after (S2) control or food patch presentations in Experiment 1. In experiment 2, focal observations were only done in between

**Figure S2.**
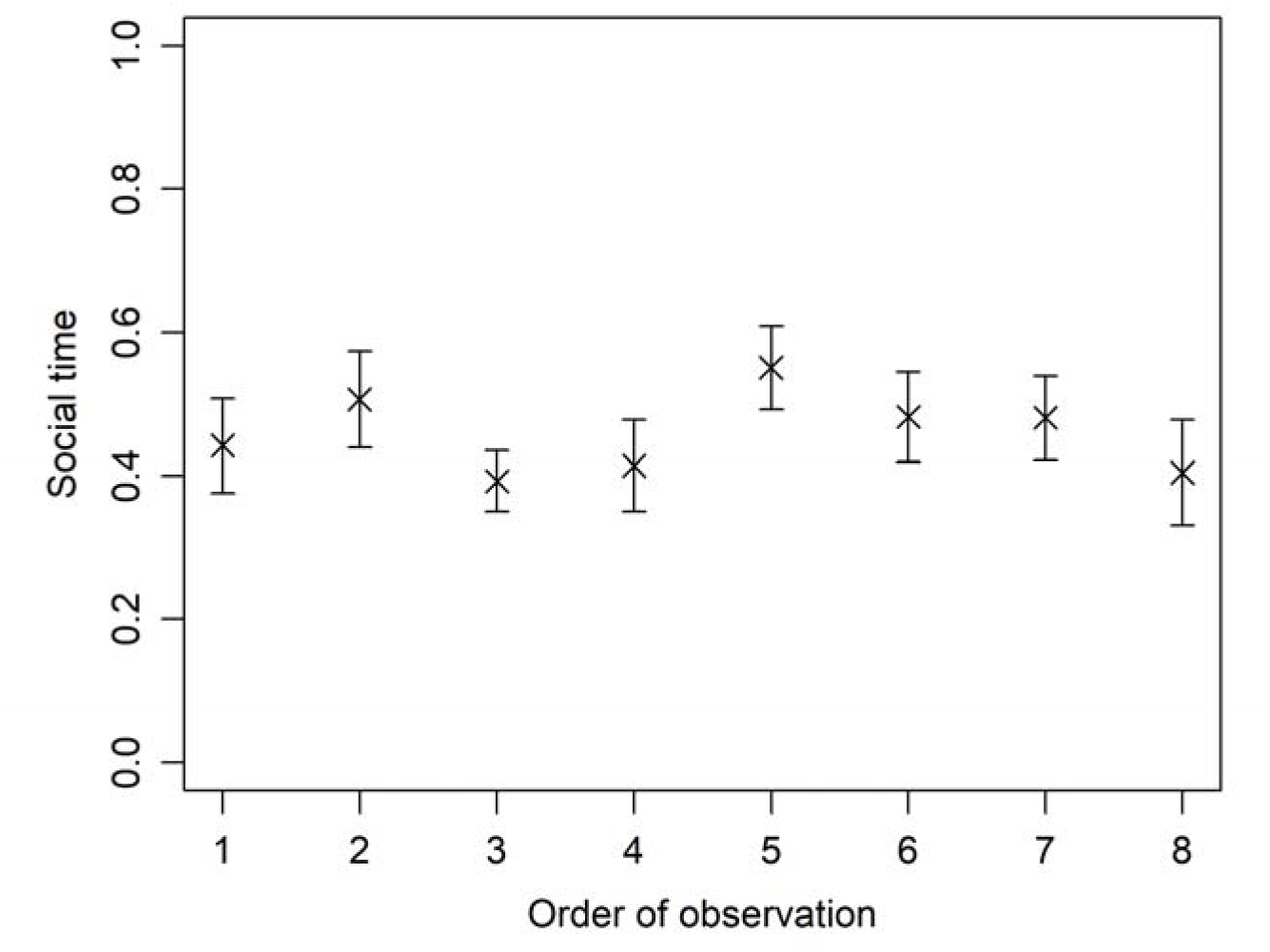
Mean individual social time ± SE in relation to order of observation after the first series of ephemeral food patches (S1).

**Figure S3.**
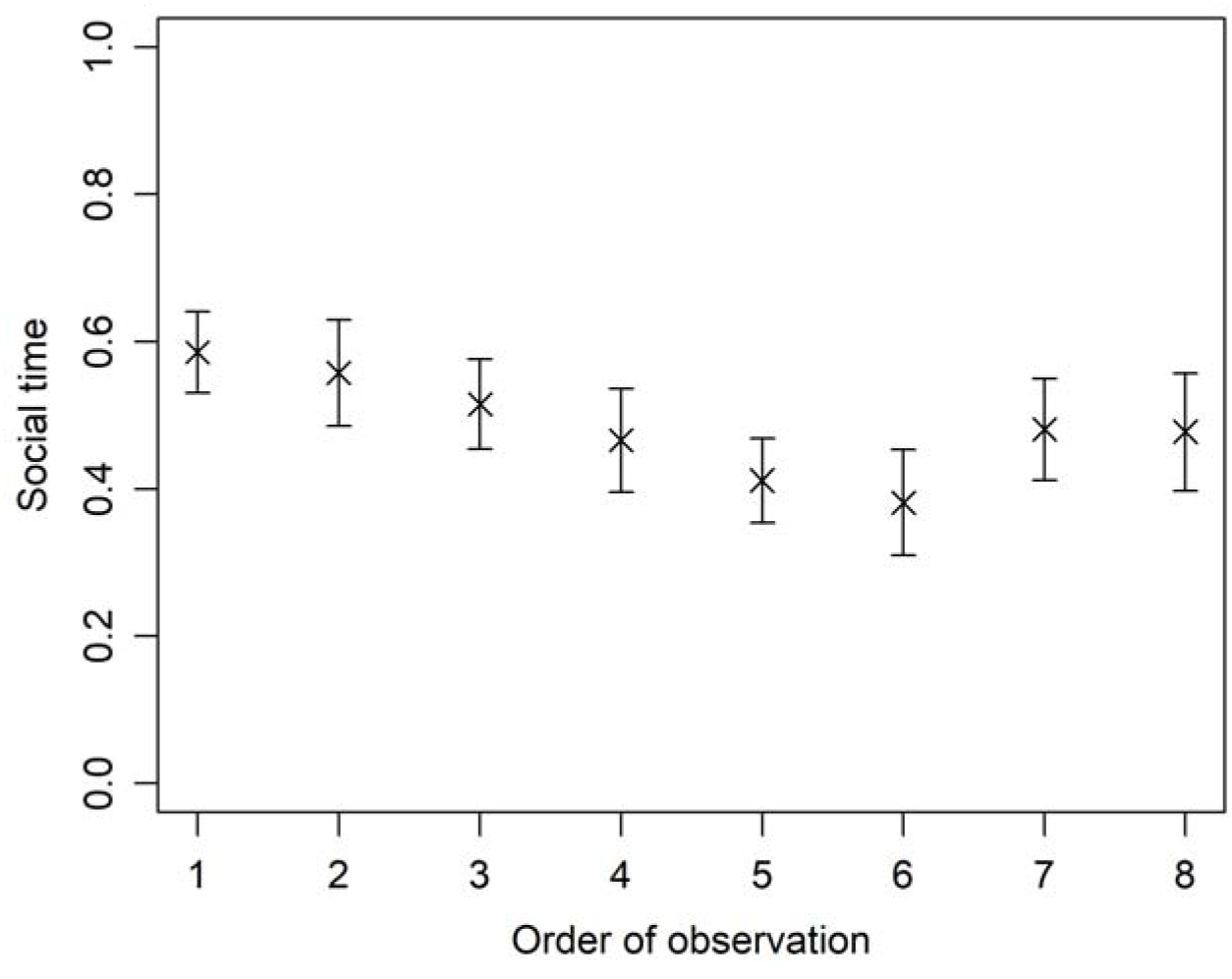
Mean individual social time ± SE in relation to order of observation after the second series of ephemeral food patches (S2).

**Figure S4.**
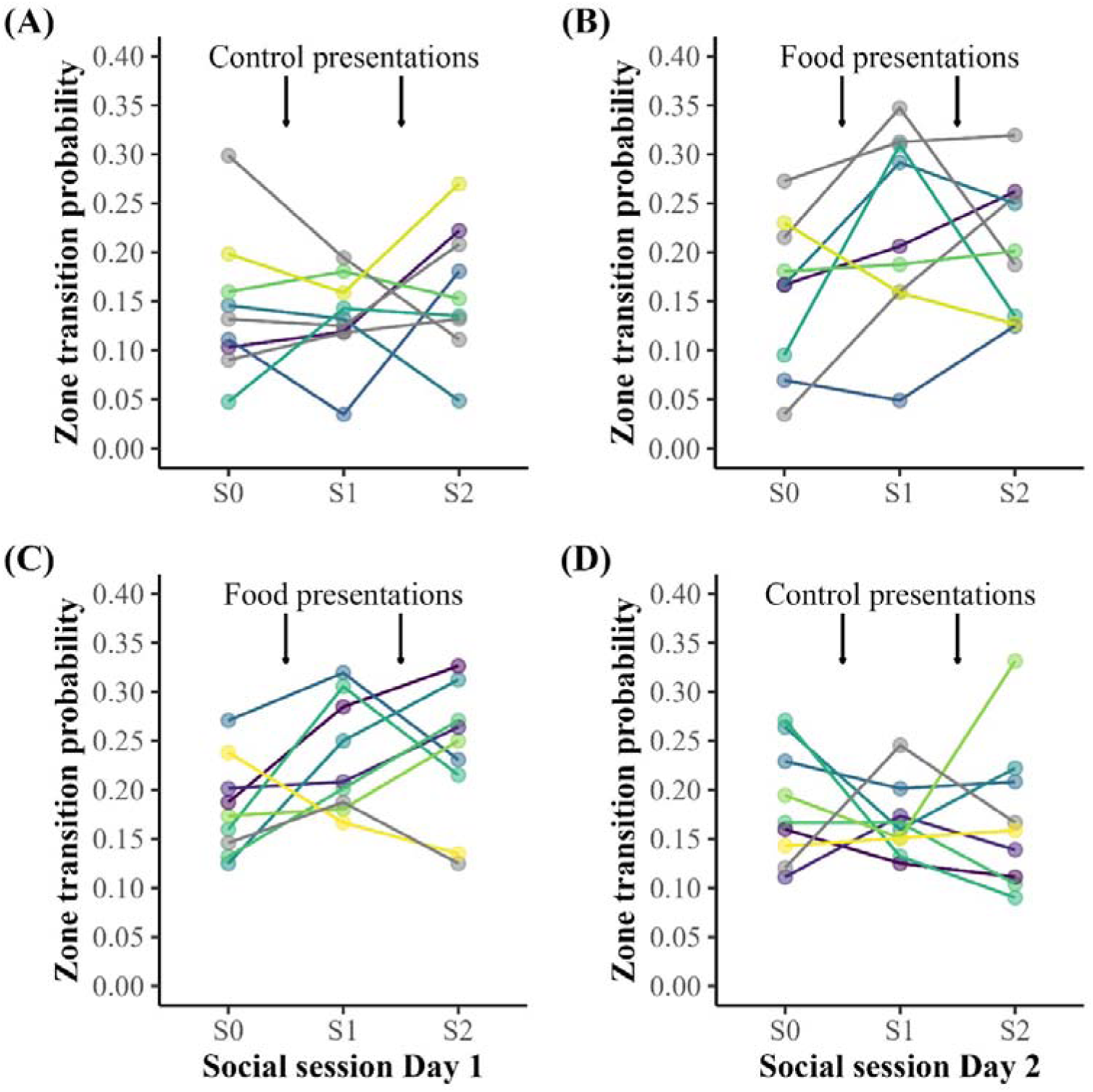
Spatial zone transition probability of wild guppy groups during social observation session before (S0), between (S1), and after (S2) two series of repeated food patch or control patch presentations. Arrows indicate the timing of the presentations. A) Groups that received the control patches on the first day B) received the food patches on the second day. C) Groups that received the food patches on the first day D) received the control patches on the second day. Pools consisted of five spatial zones, identified by the nearest patch presentation location. Colours depict group ID.

**Figure S5.**
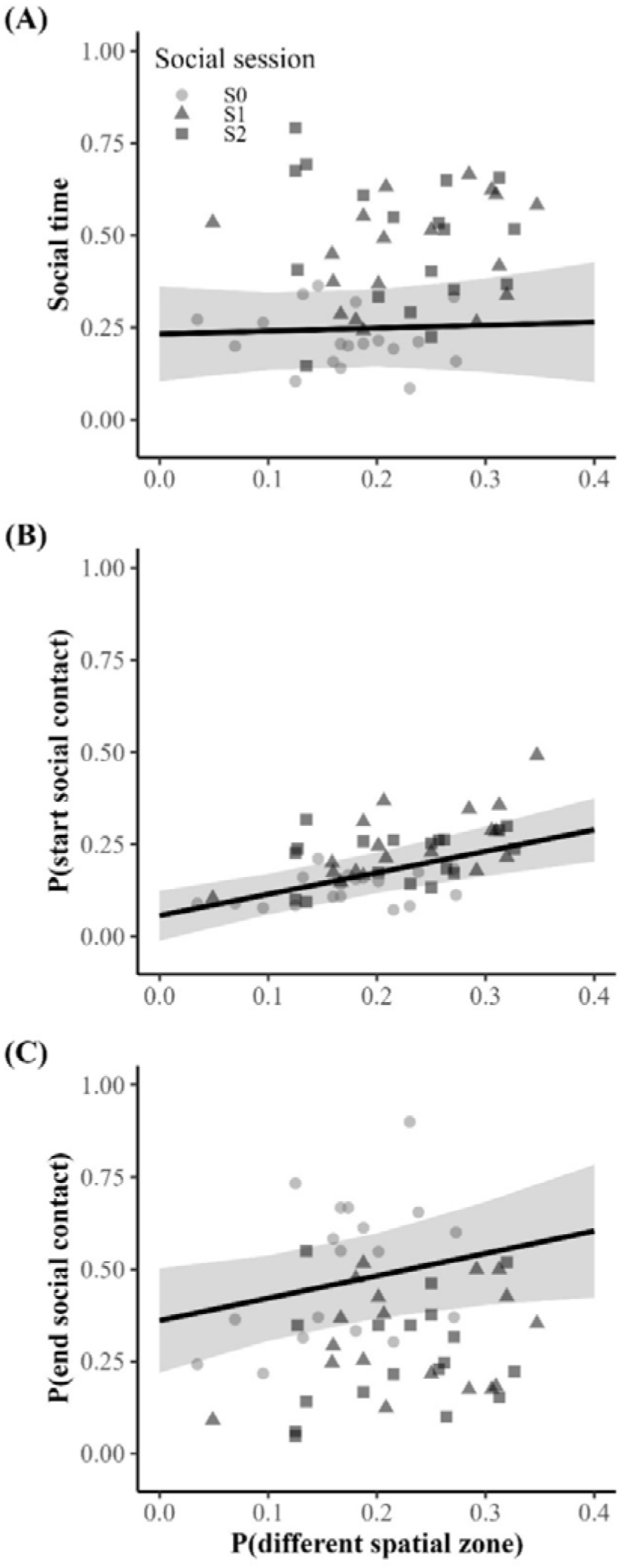
Relationships between spatial zone transition probability and the social dynamics of wild guppy groups. The x-axis represents the probability of transitioning between any of the five spatial zones in the pool (in contrast to staying in the same spatial zone) as a predictor of (A) the proportion of time spent in social contact, (B) the probability of starting social contact (Pa→s), and (C) the probability of ending social contact (Ps→a). Points represent observed data, with colour and shape indicating social session (S0–S2). Solid lines show model-predicted values from mixed-effects models, and shaded ribbons indicate 95% confidence intervals.

## Notes

https://www.osf.io/75vtn

### Competing Interest Statement

The authors have declared no competing interest.

### Summary of Updates

Most significantly, the methods and results have been restructured and rephrased to create more focus. The supporting information (SI) was extended with two tables and a figure, giving more clarity and detail about the included variables, the various models and the specific timelines of the experiments. Certain supporting analyses were removed or moved to the SI. Treatment sequence was added as a control variable to all models. None of the conclusions changed.

